# Mutual Antagonism Between PRC1 Condensates and SWI/SNF in Chromatin Regulation

**DOI:** 10.1101/2025.08.25.672128

**Authors:** Stefan Niekamp, Sharon K. Marr, Rebecca Sanon, Philipp C. Schneider, Radhika Subramanian, Robert E. Kingston

## Abstract

Opposing activities of conserved chromatin regulatory complexes, such as the Polycomb Repressive Complex 1 (PRC1) and the activating chromatin remodeler SWI/SNF play critical roles in regulating gene expression during development and differentiation. The mechanisms by which these complexes compete to regulate chromatin states remain poorly understood. We combine single-molecule analysis and genomic approaches in cultured cells to demonstrate that the condensate-forming properties of PRC1 play an important role in excluding SWI/SNF from chromatin. Consistently, PRC1 compositions with a higher propensity for condensate formation are more effective in preventing SWI/SNF binding. Conversely, SWI/SNF-bound chromatin significantly reduces PRC1 binding and subsequent condensate formation. Notably, SWI/SNF can suppress PRC1 condensate formation in an ATP-hydrolysis independent manner. We propose that the condensate properties of different PRC1 compositions drive mutual PRC1-SWI/SNF antagonism to properly balance these competing regulatory activities during development.

## Introduction

Maintaining precise gene expression levels is essential for defining cellular identity and preventing disease. In mammalian cells, gene activation and repression are governed by multiprotein complexes that regulate chromatin at both local and higher-order levels. Mutations in these chromatin-modifying complexes are associated with a wide range of diseases, including developmental disorders and cancers^1–7^. Among the key regulators of chromatin states are the highly conserved Polycomb-Group (PcG) and Trithorax-Group (trxG) proteins. These frequently function in opposition: PcG proteins normally establish repressive chromatin environments, while trxG proteins normally promote active, accessible chromatin^8–12^. This antagonistic relationship was first identified through genetic screens in *Drosophila*. Deletion of PcG genes has been shown to lead to Hox gene derepression, resulting in homeotic transformations, while trxG mutations counteract these effects^13,14^. Conversely, loss-of-function mutations in trxG genes have been shown to lead to insufficient Hox gene expression causing homeotic transformations^15^. These complexes have subsequently been shown to contribute to numerous human malignancies^5–7^.

Polycomb-group proteins are typically categorized into one of two major complex families: Polycomb repressive complex 1 (PRC1) and Polycomb repressive complex 2 (PRC2). PRC2 functions as a methyltransferase that catalyzes the trimethylation of histone H3 lysine 27 (H3K27me3), a hallmark of repressed chromatin. The PRC1 family is subdivided into variant PRC1 complexes, which mono-ubiquitinate histone H2A at lysine 119 (H2AK119ub), and canonical PRC1 (cPRC1) complexes, which alter chromatin structure through nonenzymatic roles by compacting chromatin and modulating long-range chromatin interactions^5,16,17^. Canonical PRC1, the focus of this study, consists of four core subunits (RING, PCGF, CBX, and PHC); the CBX subunit binds to H3K27me3 thereby connecting PRC1 to PRC2. Each of these four canonical PRC1 subunits has multiple paralogs, resulting in over 60 possible combinations^16,17^.

On the opposing side, the Trithorax-group includes ATP-dependent chromatin remodeling enzymes, such as the mammalian SWI/SNF (SWItch/Sucrose Non-Fermentable) complexes — also known as the BAF (BRG1/BRM-associated factor) complexes^4,18,19^. The mammalian SWI/SNF family of complexes are large, multimeric assemblies with approximately 15 subunits, many of which have multiple paralogs, giving rise to hundreds of potential combinations^20^. Together with other chromatin remodelers, SWI/SNF can generate accessibility of nucleosomal DNA, can slide nucleosomes, and can evict nucleosomes^21–23^.

The formation of biomolecular condensates has emerged as a mechanism for organizing macromolecules within the nucleus^24–31^. Both SWI/SNF^32–36^ and PRC1^37–41^ subunits as well as chromatin^42,43^ itself have been shown to form condensates *in vitro* and in cells. For instance, the SWI/SNF subunit ARID1A forms condensates that are critical for proper genomic targeting of SWI/SNF^33,34^. Interestingly, SWI/SNF can not only form condensates itself, but also de-condensate chromatin and drive micro-scale movements of chromatin condensates^44^. For canonical PRC1, the CBX and PHC subunits are thought to be the main drivers of condensate formation^37–41^. Disruption of canonical PRC1 condensate formation has been linked to impaired long-range chromatin interactions, developmental defects, and reduced gene repression in mice^45–50^ supporting the idea that PRC1 condensates are important for maintaining transcriptional silencing^10,16,37,38^. Our recent work has demonstrated that the composition of canonical PRC1 influences the biophysical properties of these condensates^40^. One prediction that arises from these studies is that different combinations of PRC1 subunits may modulate condensate dynamics and material properties, ultimately influencing PRC1’s ability to antagonize transcriptional activators such as SWI/SNF.

In this study, we build upon these findings to investigate the molecular mechanisms by which PRC1 and SWI/SNF complexes compete to regulate chromatin states. Although genetic, biochemical, and gene-targeting approaches have provided significant insights into the antagonism between Polycomb and SWI/SNF^51–55^, the collective activities of their subunits — and how these dictate the outcome of their competition — remain poorly understood. One major challenge in addressing this question is the high degree of genetic redundancy within both complexes, which complicates efforts to dissect their functions in cellular systems alone. To address these limitations, we developed a reconstitution-based single-molecule assay that enables systematic investigation of the biophysical principles governing this competition. This assay, combined with live-cell imaging and genomic analyses, allowed us to directly examine how PRC1 and SWI/SNF influence each other’s chromatin binding and activity. We found that the composition of PRC1, and the associated ability of specific subunit combinations to form condensates, plays a key role in excluding SWI/SNF from chromatin, thereby maintaining a transcriptionally silent state. Conversely, we observed that pre-bound SWI/SNF can inhibit PRC1 recruitment and prevent condensate formation through mechanisms that do not require ATP hydrolysis, indicating that competitive chromatin binding, rather than active remodeling alone, underlies this antagonistic interaction. Together, our findings reveal a mutually antagonistic relationship between PRC1 condensates and SWI/SNF underscoring the importance of PRC1 composition and condensate dynamics in regulating gene repression.

## Results

### PRC1 condensates and SWI/SNF are anti-correlated in HCT116 cells

We first examined how PRC1 and SWI/SNF interact in cells using live-cell and super-resolution imaging. For these experiments, we used HCT116 cells, a well-established cell line for SWI/SNF research^56–58^ that also expresses high levels of condensate-forming PRC1 subunits such as PHC2^39–41^ (**Figure S1A-D**). This cell line is therefore particularly well-suited to investigate the antagonism between PRC1 and SWI/SNF.

Immunofluorescence microscopy of PHC2 and Brg1 (a SWI/SNF subunit) and PHC2 and RING1B (a PRC1 subunit) in HCT116 cells revealed an anti-correlated pattern between PHC2 and Brg1, yielding Pearson correlation coefficients of 0.18 (PHC2/Brg1) and 0.65 (PHC2/RING1B) (**Figure 1A-C**). This spatial anti-correlation of PRC1 and SWI/SNF was further confirmed by live-cell confocal microscopy of endogenously HALO^59^-tagged PHC2 and mEOS3.2^60^-tagged ARID1A (**Figure S1E, F**). The enhanced signal from immunofluorescence also enabled us to perform Structured Illumination Microscopy (SIM), which provided higher-resolution imaging, making the anti-correlation between PHC2 and ARID1A even more apparent (**Figure 1D, E, S1G, Movies S1, S2**). We frequently observed ARID1A at the periphery of PHC2 condensates in cells, with minimal mixing between the two. Beyond imaging-based approaches, this anti-correlation was also evident in genomic approaches looking at chromatin binding using CUT&RUN^61^ in HCT116 cells. Only 0.3% of PHC2 peaks overlapped with BAF155 (a SWI/SNF subunit), whereas 92.1% of PHC2 peaks overlapped with RING1B (**Figure 1F**). Collectively, these findings demonstrate that PRC1 condensates and SWI/SNF show mutually exclusive chromatin binding patterns in HCT116 cells. These data led us to further explore the hypothesis that PRC1 condensates might have distinctive properties required for exclusion of SWI/SNF from repressed chromatin states.

**Figure 1.**
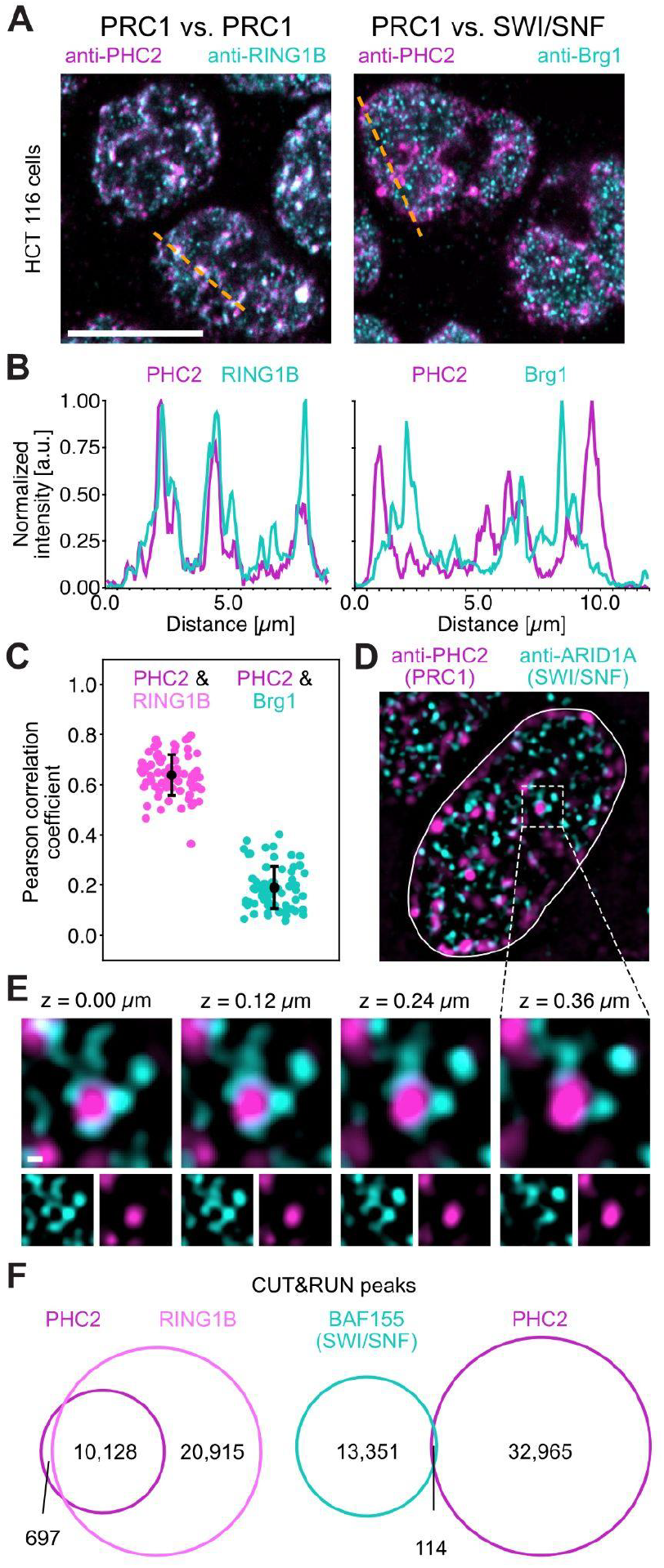
PRC1 condensates and SWI/SNF are anti-correlated in HCT116 cells. (**A**) Immunofluorescence images of HCT116 cells. The scale bar is 10 μm. (**B**) Line scan of intensities of data shown in **A** along the orange dashed lines. (**C**) Pearson correlation coefficient of intensity distributions in the nucleus of data as shown in **A**. 139 nuclei were analyzed. (**D**) Structured Illumination Microscopy (SIM) micrographs of fixed HCT116 cells. The solid white line shows the outline of the nucleus. Square with a dashed line is magnified in **E**. Data is representative of three biological replicates. (**E**) Z-slices of magnified area from data as shown in **D**. Scale bar is 200 nm. (**F**) Venn diagrams of overlapping CUT&RUN peaks in HCT116 cells (log2 > 2; adj. P < 0.0001).

### PRC1 condensates inhibit SWI/SNF binding

We investigated the role of condensate formation in the competition between PRC1 and SWI/SNF using our previously published single-molecule Total Internal Reflection Fluorescence (TIRF) microscopy protocol^40^ (**Figure 2A**). Fluorescently labeled 12-mer nucleosomal arrays, tethered to a lipid bilayer via biotinylated DNA and streptavidin, were used to observe binding of recombinant fluorescently-labeled PRC1 and SWI/SNF complexes (**Figure 2A, S2A-H**). We initially focused on mammalian SWI/SNF (cBAF) and PRC1 composed of CBX2, PHC2, RING1B, and PCGF4 (**Figure S2A, B**). When we simultaneously combined all three components, PRC1, SWI/SNF, and nucleosomal arrays, and added them to the microscopy chamber, we observed differential binding for PRC1 and SWI/SNF to nucleosomal arrays (**Figure 2B, C, Movie S3**). Specifically, we found that SWI/SNF occupies single nucleosomal arrays while PRC1 is predominantly observed in condensates with tens of nucleosomal arrays.

**Figure 2.**
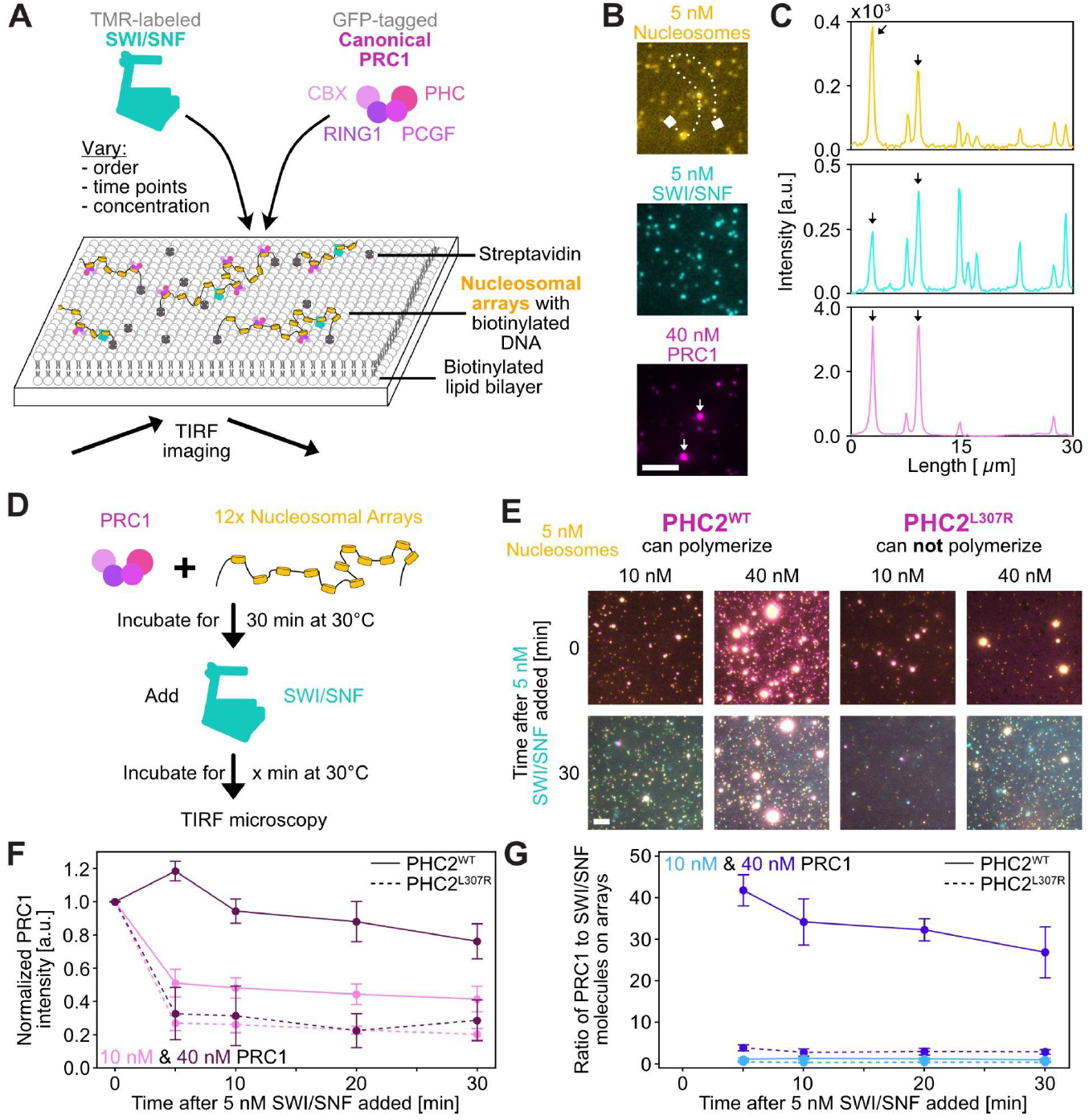
PRC1 condensates inhibit SWI/SNF binding. (**A**) Work-flow of Total Internal Reflection Fluorescence (TIRF) microscopy assay to study competition of PRC1 and SWI/SNF for access to nucleosomal arrays (see **Methods** for details). (**B**) Representative TIRF micrographs of nucleosomal arrays (yellow), SWI/SNF (cyan), and PRC1 (pink) 30 minutes after all were mixed simultaneously. The scale bar is 5 μm. (**C**) Line scan of intensities of the images in **B** along the dotted line. (**D**) Schematics showing the order of mixing for nucleosomal arrays, SWI/SNF, and PRC1. (**E**) Example TIRF micrographs of nucleosomal arrays, SWI/SNF, and PRC1 with PHC2^WT^ or PHC2^L307R^. The scale bar is 10 μm. (**F**) Change of average PRC1 intensities on nucleosomal arrays over time. (**G**) Ratio of PRC1 to SWI/SNF molecules bound to nucleosomal arrays over time. (**F, G**) Error bars are standard error of the mean of four technical replicates. For each condition 12,000 to >130,000 spots have been analyzed.

Based on this initial observation, we hypothesized that if PRC1 condensates determine how well SWI/SNF can access chromatin, we should see differences in SWI/SNF binding to nucleosomal arrays in the presence of PRC1 complexes with distinct condensate properties. To test this, we focused on the PHC subunit, known for its role in PRC1 condensate formation^39–41^. The PHC subunit’s C-terminal sterile alpha motif (SAM) domain facilitates head-to-tail polymerization^62^, a process linked to long-range chromatin interactions and gene repression^46,49,50^. Moreover, the polymerization deficient mutant PHC2 L307R (PHC2^L307R^) has been shown to form liquid-like condensates compared to the gel-like condensates formed by PRC1 with PHC2 wild type (PHC2^WT^)^40^. We first investigated PRC1 containing PHC2^WT^ and pre-incubated PRC1 and nucleosomal arrays for 30 minutes and subsequently added SWI/SNF (**Figure 2D**). We used PRC1 concentrations of 10 nM and 40 nM, which represent low and high degrees of condensate formation, respectively (9% vs. 80% of PRC1 molecules in condensates; **Figure S2I**). At 10 nM PRC1 we found that 52% of PRC1 was evicted 10 minutes after the addition of 5 nM SWI/SNF, while only 6% of PRC1 was displaced at 40 nM PRC1 (**Figure 2E-G**). We further verified these observations by comparing the competition across a broad range of PRC1 and SWI/SNF concentrations and found that SWI/SNF could only bind effectively to arrays at PRC1 concentrations below 10 nM, the threshold for PRC1 condensate formation (**Figure S2J-L**). In addition, we used an orthogonal epifluorescence microscopy approach under conditions that promote larger PRC1 condensates and observed that added SWI/SNF preferentially accumulates at the droplet periphery, with a fourfold higher intensity than at the droplet center (**Figure S2M–O**). These findings are consistent with the cellular super-resolution imaging data (**Fig. 1D, E**). Together, these results suggest a mechanism in which PRC1 condensates exclude SWI/SNF from repressed chromatin states.

We asked whether the different condensate properties of PRC1 containing PHC2^WT^ versus PHC2^L307R^ could change the outcome of the competition between PRC1 and SWI/SNF using the single-molecule TIRF assay. Interestingly, our comparison of PRC1 with PHC2^WT^ versus PHC2^L307R^ at the same concentrations demonstrated a substantial functional difference in their ability to oppose SWI/SNF, with PHC2^L307R^ being far less effective (**Figure 2E-G, S3A, B**). For example, in an experiment with 5 nM SWI/SNF and 40 nM PRC1, only 6% of PRC1 was displaced from nucleosomal arrays after 10 minutes of SWI/SNF addition for PRC1 with PHC2^WT^ while 69% of PRC1 was evicted for PRC1 with PHC2^L307R^. Moreover, we found that PRC1 with PHC2^WT^ under condensate forming concentrations was able to maintain a more than 25-fold higher ratio of PRC1 over SWI/SNF in binding to nucleosomal arrays, while for PRC1 with PHC2^L307R^ the ratio was approximately 1:1 (**Figure 2G**). These observations were further supported by an orthogonal epifluorescence microscopy approach (**Figure S3C-H**). We conclude that altering the nature of PRC1 condensate through the use of PHC2 mutation or through the titration of PRC1 concentrations impacts the ability of PRC1 to restrict SWI/SNF’s access to nucleosomes.

### PRC1 composition dictates outcome of competition with SWI/SNF

We further investigated the role of PRC1 condensates in opposing SWI/SNF by examining complexes with different CBX subunits, which are known to alter the degree PRC1 condensate formation^37,38,63,40^. Specifically, we focused on the paralogs CBX2 (CBX2^WT^) and CBX7 (CBX7^WT^), where CBX2 displays strong nucleosomal array compaction and condensate formation while CBX7 does not share these properties at the same con- centration^45,38,40^. We also characterized two CBX2 mutants that have moderate (CBX2^13KRA^) or severe (CBX2^23KRA^) defects in condensate formation^38,40,64^ (**Figure 3A, B**). We found that SWI/SNF competed more effectively with the complexes with lower ability to form condensates (CBX7^WT^ and CBX2^23KRA^ containing complexes) than it did with CBX2^WT^ and the less defective CBX2^13KRA^ containing complexes (**Figure 3C**). When 20 nM SWI/SNF was added to 20 nM PRC1 with CBX2^WT^ the PRC1 intensity only dropped to 27% compared to when no SWI/SNF was added while it decreased to 3% for PRC1 with CBX7^WT^ (**Figure 3C**). Moreover, the ratio of PRC1 over SWI/SNF molecules bound to nucleosomal arrays is more than 4-fold higher for PRC1 with CBX2^WT^ compared to PRC1 with CBX7^WT^ (**Figure 3D, S3I**). Thus, these data highlight a correlation between the ability of PRC1 to form condensates and its ability to block SWI/SNF from accessing nucleosomes. Together with our observations for the PHC2 subunit, we conclude that the PRC1 composition influences the degree of PRC1 eviction by SWI/SNF.

**Figure 3.**
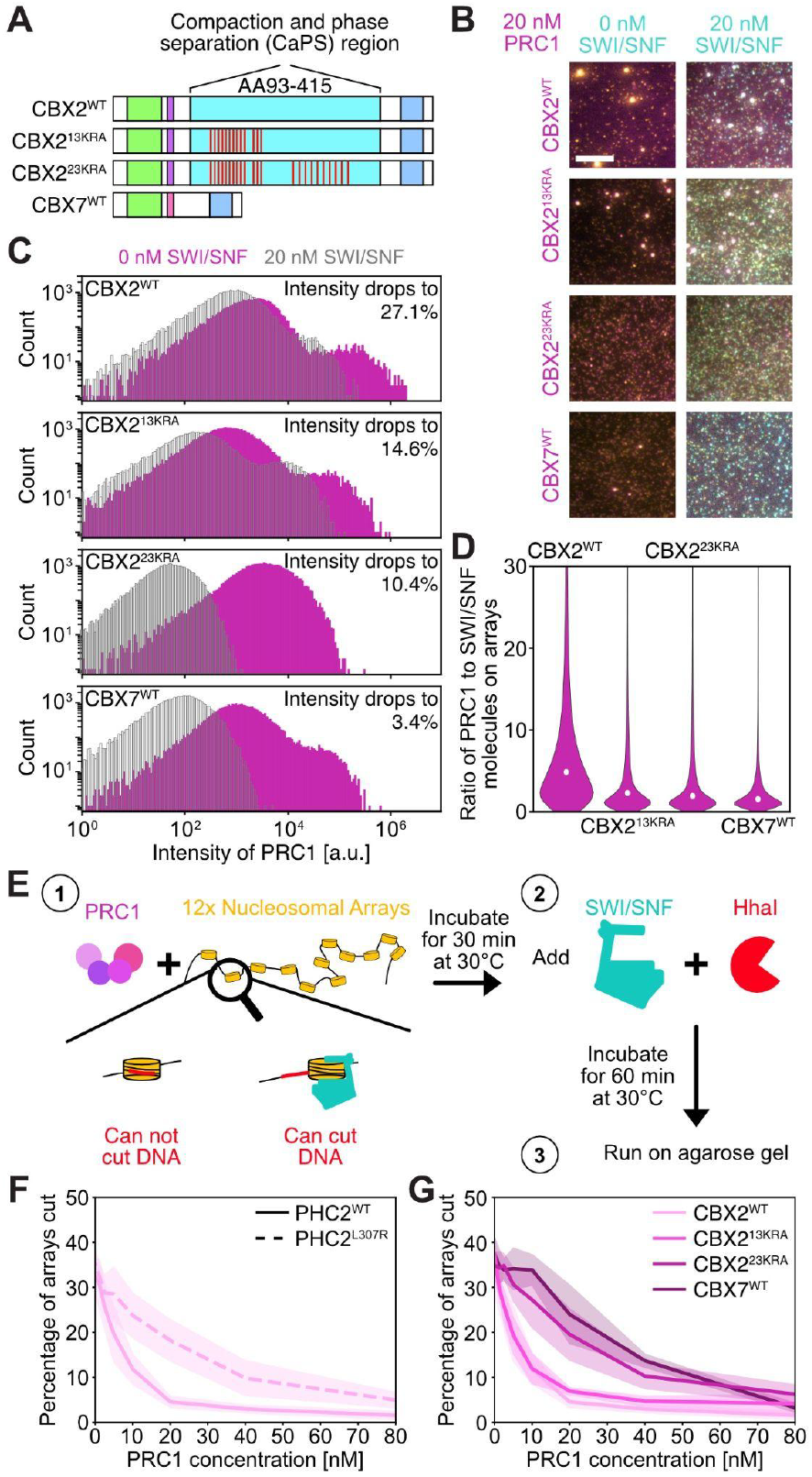
PRC1 composition dictates the outcome of competition with SWI/SNF. (**A**) Schematics of different CBX constructs. (**B**) Representative TIRF micrographs of nucleosomal arrays (yellow), SWI/SNF (cyan), and PRC1 (pink) with different CBX subunits. The scale bar is 10 μm. (**C**) Intensity histograms of PRC1 with different CBX subunits with 0 or 20 nM SWI/SNF. (**D**) Ratio of PRC1 to SWI/SNF molecules bound to nucleosomal arrays of PRC1 with different CBX subunits in the presence of 20 nM SWI/SNF. (**C, D**) For each condition 33,000 to >59,000 spots were analyzed. Data are representative of at least two technical replicates. (**E**) Schematic of Restriction Enzyme Accessibility (REA) assay. (**F, G**) Percentage of nucleosomal array cutting in REA assay for different PRC1 compositions. Opaque area is standard error of the mean of three replicates.

To test whether the reduction in SWI/SNF binding to nucleosomal arrays by PRC1 condensates also leads to a reduction in chromatin remodeling, we employed a Restriction Enzyme Accessibility (REA) assay (**Figure 3E**). Similarly to the TIRF assay, we allowed PRC1 to interact with nucleosomal arrays for 30 minutes before adding SWI/SNF and the HhaI restriction enzyme. The HhaI enzyme can only cut DNA when the center nucleosome has been moved or removed by SWI/SNF remodeling activity. Thus, cut DNA will only be visible on an agarose gel, if SWI/SNF can remodel nucleosomes. In agreement with previous work^38,45^ and our imaging data, we found that SWI/SNF chromatin remodeling is more inhibited for PRC1 compositions that form larger condensates at the same concentration (**Figure 3F, G**). For instance, at 20 nM PRC1, only 4% of the DNA is cut for PRC1 with CBX2^WT^ while 24% of DNA is cut for PRC1 with CBX7^WT^. Consistently, the CBX2^13KRA^ containing complex blocked remodeling to a lower extent relative to the CBX2^23KRA^ containing complex, which is consistent with the impact of these mutations on condensate formation. Taken together, we conclude that PRC1 composition and the resulting condensate properties dictate the outcome of the competition between PRC1 and SWI/SNF for binding to nucleosomal arrays and thereby modulate SWI/SNF’s ability to remodel chromatin.

### SWI/SNF occupancy on chromatin restricts PRC1 binding and condensate formation

Nucleosomal arrays and PRC1 act synergistically to form condensates and full PRC1 occupancy of nucleosomal arrays results in a reduction of the critical concentration required for condensation by more than 20-fold^40^. We thus asked what happens to PRC1 condensate formation and binding if nucleosomal arrays are pre-bound by SWI/SNF. To test this, we pre-incubated nucleosome arrays with SWI/SNF and added PRC1 after 30 minutes (**Figure 4A**). We found that PRC1 binding at condensate forming concentrations (e.g 40 nM) is significantly reduced when SWI/SNF is already present on nucleosomal arrays (**Figure 4B-D, S4A-H**). For instance, the PRC1 intensity at 40 nM PRC1 is 3-fold lower when 20 nM SWI/SNF is added first compared to when PRC1 is present first (**Figure 4B, C**). This reduction in binding of PRC1 to nucleosomal arrays also resulted in a significant reduction of PRC1 condensate formation. For instance, when 20 nM SWI/SNF are pre-incubated with nucleosomal arrays only 5% of PRC1 molecules are in condensates at 40 nM PRC1 concentration while 78% of PRC1 molecules are in condensates when PRC1 is added first (**Figure 4D**). In contrast, at concentrations at which PRC1 does not form condensates even in the absence of SWI/SNF (e.g. 2.5 nM), we found no difference in terms of PRC1 binding to nucleosomal arrays (**Figure S4A, B**). This data is in good agreement with our earlier observations (**Figure 2 and 3**) showing that PRC1 in condensates are resistant to eviction by SWI/SNF.

**Figure 4.**
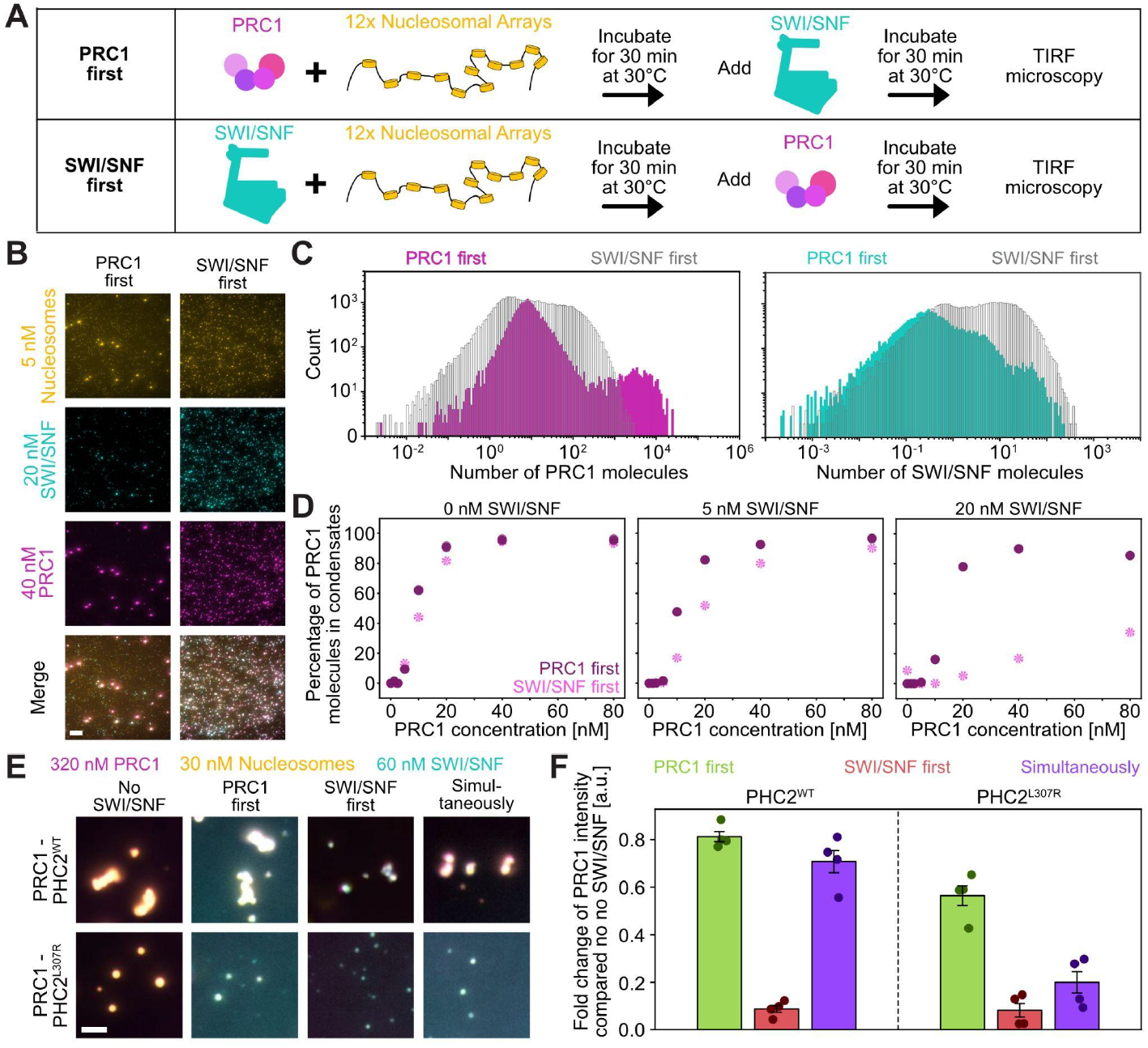
SWI/SNF occupancy on chromatin restricts PRC1 binding and condensate formation. (**A**) Schematics showing the order of mixing for nucleosomal arrays, SWI/SNF, and PRC1. (**B**) Representative TIRF micrographs when SWI/SNF (cyan) and PRC1 (pink) are added to nucleosomal arrays (yellow) in different orders. The scale bar is 10 μm. (**C**) Intensity histograms of PRC1 and SWI/SNF for different order of addition. (**D**) Percentage of PRC1 molecules in condensates when SWI/SNF (pink) or PRC1 (purple) were added first to nucleosomal arrays. (**C, D**) For each condition 29,000 to >105,000 spots were analyzed. Data are representative of at least two technical replicates. (**E**) Representative epifluorescence microscopy micrographs when SWI/SNF and PRC1 are added to nucleosomal arrays in different orders for PRC1 complexes with PHC2^WT^ or PHC2^L307R^. The scale bar is 5 μm. (**F**) Bar graphs of PRC1 intensity change for data as shown in **E**. Error bars are standard error of the mean of four technical replicates. For each condition 1,400 to >11,000 spots were analyzed.

We further validated these findings utilizing a common droplet assay in which condensates were imaged by epifluorescence microscopy. We compared how PRC1 with PHC2^WT^ or PHC2^L307R^ fare when PRC1 or SWI/SNF are added first to nucleosomal arrays or when all three are mixed simultaneously (**Figure 4E**). In agreement with our single-molecule observations (**Figure 4B-D**), we found that adding SWI/SNF first greatly reduces PRC1 binding and condensate formation compared to when PRC1 is present first on nucleosomal arrays. Interestingly, we observed that when all three components are mixed simultaneously, that PRC1 with PHC2^WT^ forms condensates similar to those formed when PRC1 is added first, while PRC1 with PHC2^L307R^ forms condensates similar to when SWI/SNF is added first (**Figure 4F**). This again highlights the role of PRC1 condensate properties in modulating the competition with SWI/SNF. Taken together, we conclude that SWI/SNF occupancy on chromatin can restrict PRC1 binding and condensate formation.

### SWI/SNF suppresses PRC1 condensate formation in an ATP-hydrolysis independent manner

One of SWI/SNF’s important roles is to remodel chromatin in an ATP-dependent manner by altering nucleosome structure and/or occupancy^21^. We thus asked whether ATP is important for SWI/SNF to compete with PRC1 for access to nucleosomes and made two different ATPase mutants of SWI/SNF in the Brg1 subunit. We used a previously characterized Walker A mutant^53,65–71^ (SWI/SNF^K785R^) which is implicated in preventing ATP binding and a Walker B mutant^53,69,70^ (SWI/SNF^D881N^) which is implicated in preventing ATP hydrolysis (**Figure 5A, S5A**). We first tested these mutant remodelers and wild type SWI/SNF (SWI/SNF^WT^) in the REA assay and found - as expected - no nucleosome remodeling for the ATPase mutants (**Figure 5B**). We next performed TIRF assays where we pre-incubated PRC1 and nucleosomal arrays, and subsequently added SWI/SNF^WT^ or the mutants. We found that both ATPase mutants bound more strongly to nucleosomal arrays and were better at evicting PRC1 compared to SWI/SNF^WT^ (**Figure 5C-F, S5B, C**). A previously observed increase in nucleosome binding time of these ATPase mutants^70–72^ likely explains their superior competitive ability against PRC1. We hypothesized that these ATPase-deficient remodelers should be particularly effective when they establish occupancy on nucleosomal arrays prior to PRC1 addition. Consistent with this hypothesis, pre-incubation of nucleosomal arrays with 10 nM SWI/SNF^K785R^ completely blocked PRC1 condensate formation (0%) upon subsequent addition of 20 nM PRC1, whereas 10 nM SWI/SNF^WT^ pre-incubation only reduced it to 10% (**Figure S5D, E**). Performing these experiments with a non-hydrolysable ATP analog AMPPNP using the droplet assay further confirmed that the reduction in ATP cycling of SWI/SNF and the resulting longer dwell-times on nucleosomes^70–72^ results in reduced PRC1 access to chromatin (**Figure S5F, G**). Thus, we conclude that SWI/SNF can suppress PRC1 condensate formation in an ATP-hydrolysis independent manner suggesting a mechanism driven by competitive binding and steric hindrance rather than active remodeling alone.

**Figure 5.**
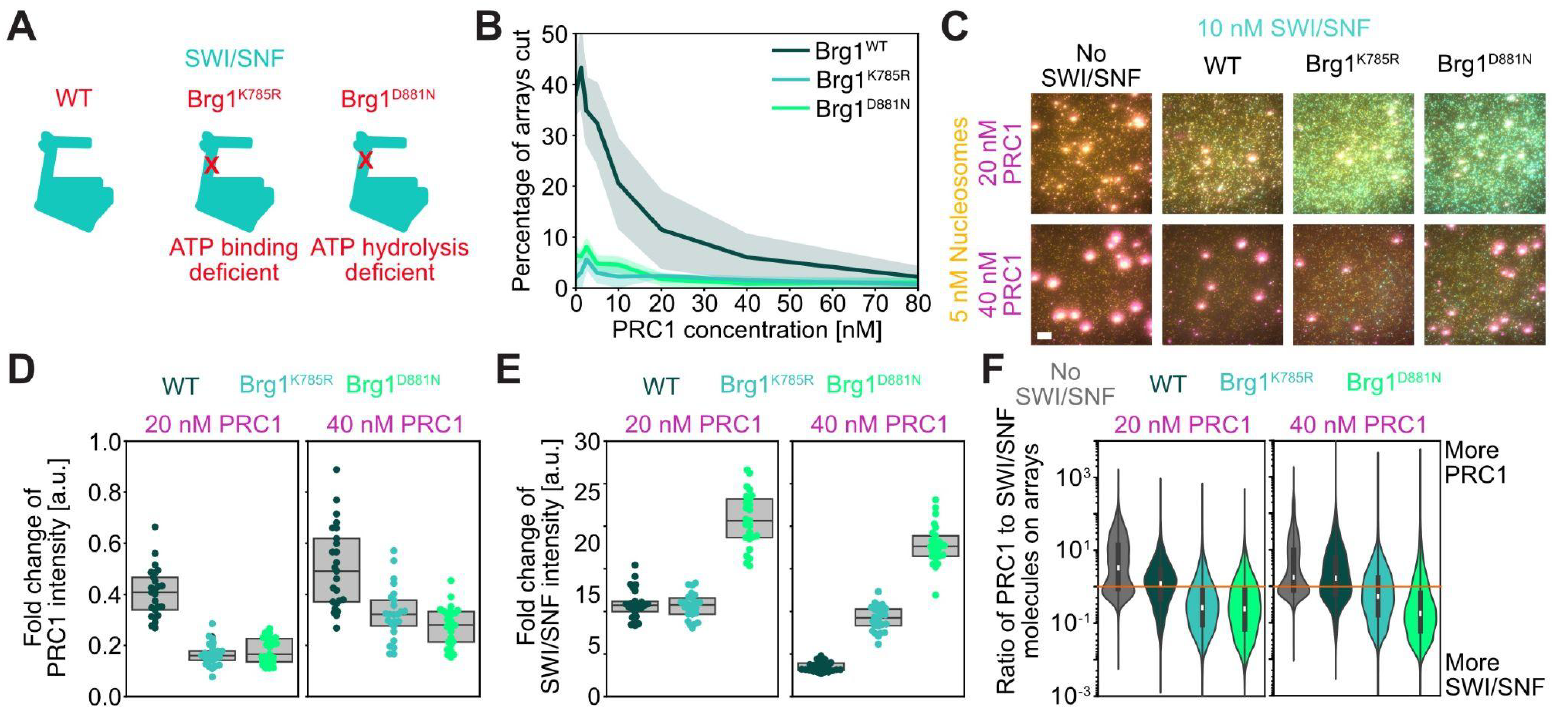
SWI/SNF suppresses PRC1 condensate formation in an ATP-hydrolysis independent manner. (**A**) Cartoon of SWI/SNF with different ATPase mutations. (**B**) Percentage of nucleosomal array cutting in REA assay for different SWI/SNF complexes. Opaque area is standard error of the mean of three replicates. (**C**) Representative TIRF micrographs of nucleosomal arrays, PRC1, and different SWI/SNF complexes when PRC1 is added first. The scale bar is 10 μm. (**D, E**) Box plot of fold change of (**D**) PRC1 and (**E**) SWI/SNF intensity. (**F**) Violin plots of the ratio of PRC1 to SWI/SNF molecules bound to nucleosomal arrays. The orange line indicates when SWI/SNF and PRC1 molecules are present in equal numbers on nucleosomal arrays. (**D-F**) Data are representative of at least two technical replicates. For each condition 44,000 to >98,000 spots were analyzed.

### Loss of PRC1 condensates leads to increased SWI/SNF binding in cells

The *in vitro* reconstitution experiments demonstrate that exclusion of SWI/SNF from chromatin increases with increasing formation of condensates. To test whether PRC1 condensate properties are important in excluding SWI/SNF from repressed genes in mammalian cells, we generated HCT116 cells with endogenous polymerization deficient PHC2 L307R mutants (PHC2^L307R^) (**Figure 6A**). This polymerization deficient PHC2 expressed at similar levels as wild type PHC2 (PHC2^WT^) (**Figure 6B**). As reported in previous studies^40,46^, PHC2^L307R^ also resulted in a significant reduction of PHC2 and RING1B condensates in HCT116 cells (**Figure 6C**). Here, we found on average zero puncta per nucleus for cells with PHC2^L307R^ while cells with PHC2^WT^ had seven puncta per nucleus (**Figure 6D**). We then compared global gene transcription levels and chromatin binding of PRC1 and SWI/SNF subunits in cells with PHC2^WT^ and PHC2^L307R^ using RNAseq and CUT&RUN^61^, respectively. We found that the polymerization deficiency of PHC2 results in a global loss of PHC2 binding (**Figure 6E-I, S6A-G**). For instance, at the transcription factor ZBTB18 locus, we observed a significant reduction in PHC2 binding which coincided with a reduction in RING1B binding, a loss in H3K27me3, and an increase in transcription (**Figure 6E**). In good agreement with work from Isono et al.^46^, we found that transcription was upregulated in the PHC2^L307R^ mutant cell line at sites that are occupied by canonical PRC1 in PHC2^WT^ cells as indicated by PHC2 and RING1B binding and H3K27me3 marks (**Figure 6F-H, S6A-D**). Of the initially more than 14,000 PHC2 peaks in wild type cells only 18 remained in the PHC2^L307R^ mutant cell line (**Figure 6H**). Interestingly, at some sites where PCH2 binding was lost as a result of the polymerization deficient mutant, we noticed an increase in SWI/SNF binding as evident from increased CUT&RUN^61^ signal (**Figure 6E, I, J, S6E-H**). For instance, at around 40 loci at which PHC2 binding was significantly decreased in PHC2^L307R^ cells, BAF155 binding increased significantly (**Figure 6J, S6H**). This increase in SWI/SNF coincided with an increase in the active histone mark H3K27ac for some genes (**Figure 6E, S6E-H**). The loss in PHC2 binding and the gain in BAF155 binding in PHC2^L307R^ cells compared to PHC2^WT^ cells often correlated with an increase in transcription suggesting that the recruitment of SWI/SNF enabled expression of these genes (**Figure 6K**). Taken together, these data are consistent with the hypothesis that PCH2 polymerization and PRC1 condensates are important to restrict SWI/SNF binding to chromatin, thereby maintaining a transcriptionally silent state.

**Figure 6.**
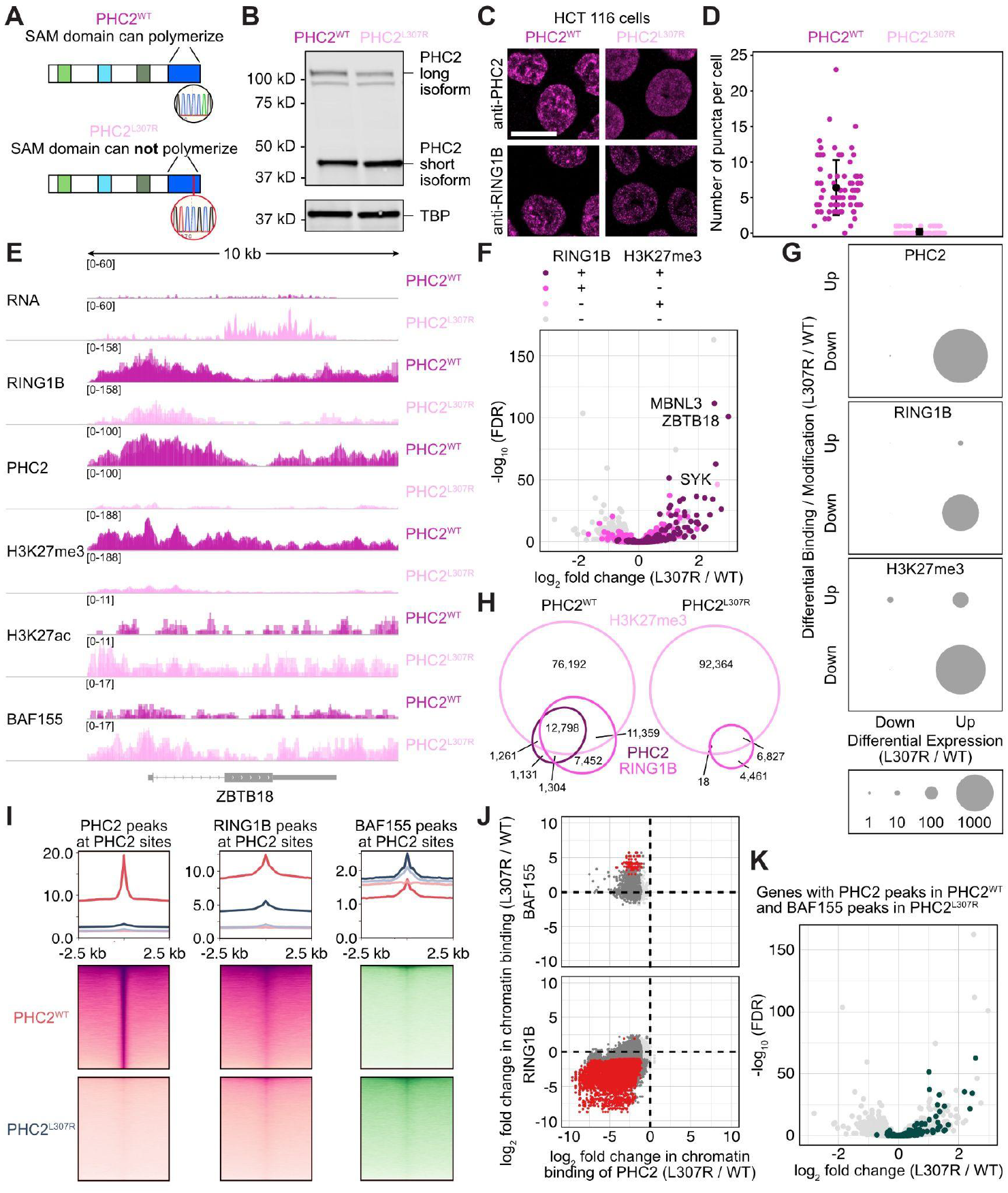
Loss of PRC1 condensates leads to increased SWI/SNF binding and transcription. (**A**) Schematics of different PHC2 constructs used to make endogenous mutations in HCT116 cells. (**B**) Immunoblot of HCT116 cells expressing endogenous PHC2^WT^ or PHC2^L307R^. Antisera against TATA-binding protein (TBP) was used as loading control. (**C**) Immunofluorescence images of HCT116 cells expressing endogenous PHC2^WT^ or PHC2^L307R^. The scale bar is 10 μm. (**D**) Quantification of the number of PHC2 puncta per nucleus of data as shown in **C**. 73 nuclei were analyzed. (**E**) Genome browser screenshots of RNAseq and CUT&RUN comparing HCT116 cells expressing endogenous PHC2^WT^ or PHC2^L307R^. (**F**) Volcano plot reflecting changes in gene expression comparing HCT116 cells expressing endogenous PHC2^WT^ or PHC2^L307R^. Pink and purple colored dots are genes reflecting changes in CUT&RUN peaks for RING1B and H3K27me3 (log2 > 1; adj. P < 0.01). (**G**) Dot plot showing number of genes with differential binding and expression patterns for PHC2, RING1B, and H3K27me3 (log2 > 1; adj. P < 0.01). (**H**) Venn diagrams of overlapping CUT&RUN peaks in HCT116 cells (log2 > 2; adj. P < 0.0001). (**I**) Heat maps showing CUT&RUN enrichment at PHC2 peak sites for different antibodies. IgG enrichment is shown as control (opaque lines, heatmaps are shown in **Figure S6G**). (**J**) Scatter plots of differential binding. Red dots are significant (log2 > 1; adj. P < 0.01) for both antibodies, dark grey for one antibody and light grey for neither antibody. (**K**) Volcano plot reflecting changes in gene expression comparing HCT116 cells expressing endogenous PHC2^WT^ or PHC2^L307R^. Dark green colored dots are genes that have significant changes in CUT&RUN peaks for both, PHC2 in PHC2^WT^ cells and for BAF155 in PHC2^L307R^ cells (log2 > 1; adj. P < 0.01). (**E-K**) All experiments were performed with four to six replicates (two different clones and two to three technical replicates each).

### SWI/SNF occupancy restricts PRC1 binding in HCT116 cells

Given that PHC2^L307R^ resulted in a global loss of PHC2 binding and an increase in SWI/SNF binding at some loci, we asked whether restoration of wild type PHC2 could reclaim sites that now had increased SWI/SNF binding. This would allow a test in cells of the *in vitro* finding that pre-binding of SWI/SNF can block PRC1 binding. To this end, we first created a frameshift mutation in PHC2 resulting in a full knockout (PHC2^KO^) of the short and long PHC2 isoforms (**Figure 7A, S7A**). We then re-expressed the dominant short isoform of PHC2 (PHC2^Re^) to wild type levels using lentiviral integration and doxycycline induction over 48 hours (**Figure 7A, S7A**). Immunofluorescence microscopy of PHC2^WT^, PHC2^KO^, and PHC2^Re^ cell lines showed a full loss of PHC2 condensates in cells with PHC2^KO^, while re-expression of PHC2 rescued condensate formation with a similar number of puncta compared to wild type cells (**Figure 7B, C**).

**Figure 7.**
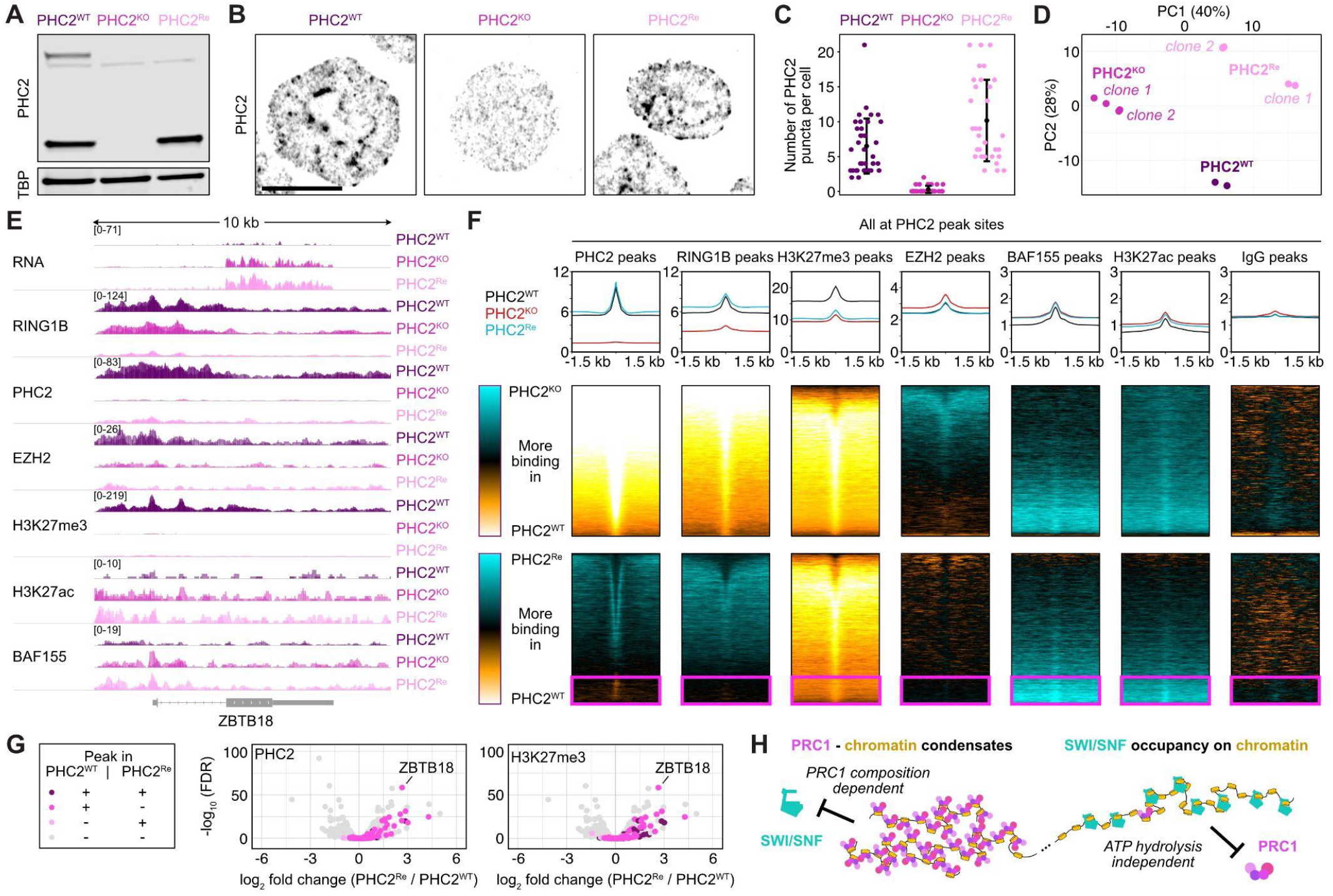
SWI/SNF occupancy restricts PRC1 binding in HCT116 cells. (**A**) Immunoblot of HCT116 cells with wild type PHC2 (PHC2^WT^), knockout of PHC2 (PHC2^KO^), and re-expressed PHC2 (PHC2^Re^). Antisera against TATA-binding protein (TBP) was used as loading control. (**B**) Immunofluorescence images of HCT116 cells with PHC2^WT^, PHC2^KO^, or PHC2^Re^. Scale bar is 10 μm. (**C**) Quantification of the number of puncta per nucleus of data as shown in **B**. 32 nuclei were analyzed. (**D**) Principal component analysis of expression profiles of HCT116 cells with PHC2^WT^, PHC2^KO^, or PHC2^Re^. (**E**) Genome browser screenshots of RNAseq and CUT&RUN comparing HCT116 cells with PHC2^WT^, PHC2^KO^, or PHC2^Re^. (**F**) Difference maps comparing CUT&RUN enrichments at PHC2 peak sites for different antibodies in HCT116 cells with PHC2^WT^ vs. PHC2^KO^ (top row) and PHC2^WT^ vs. PHC2^Re^ (bottom row). Pink box indicates loci at which PHC2 bound weaker or not at all to chromatin in the PHC2^Re^ line compared to the PHC2^WT^ line. Heat maps showing CUT&RUN enrichment for each cell line separately are shown in **Figure S7G, H**. (**G**) Volcano plot reflecting changes in gene expression comparing HCT116 cells with PHC2^WT^ or PHC2^Re^. Pink and purple colored dots are genes reflecting changes in CUT&RUN peaks for PHC2 (left) or H3K27me3 (right) in the respective cell line (log2 > 3; adj. P < 0.01). (**H**) Model for the competition of PRC1 and SWI/SNF for access to chromatin. Details are found in the **Discussion**. (**D-G**) All experiments were performed with four replicates (two different clones and two technical replicates each).

We next compared gene expression levels and chromatin occupancy across all three cell lines using RNAseq and CUT&RUN, respectively. For all three lines we found different gene expression profiles with PHC2^WT^ being closer to the PHC2^Re^ line along the main direction of change (first principal component) suggesting that the re-expression of PHC2 does not fully restore gene expression to wild type levels (**Figure 7D**). For the PHC2^KO^ line, we observed a similar global loss of PHC2 and RING1B binding and a similar de-repression as in the PHC2^L307R^ line indicating that the loss in polymerization activity results in an apparent full loss of PHC2 function (**Figure 7E, F, S7B-G**). Similarly to the PHC2^L307R^ line, SWI/SNF was also able to occupy some sites in the PHC2^KO^ line that were bound by PHC2 in wild type cells supporting the conclusion that PRC1 binding is important to restrict SWI/SNF binding to chromatin (**Figure 7E, F, S7D-H**). For the cell line with re-expressed PHC2, we found that PHC2 and RING1B can reclaim more than 90% of the sites that were lost in the PHC2 knockout line and that PHC2 and RING1B slightly exceeded binding levels compared to the wild type line (**Figure 7F, S7G, H**). However, the loci that were not re-occupied by re-expressed PHC2 to wild type levels (pink box in **Figure 7F**) were mostly loci that were bound more strongly by SWI/SNF when PHC2 was knocked out (**Figure 7E, F, S7B**). At these loci SWI/SNF remained bound at higher levels compared to wild type even when PHC2 was re-expressed suggesting that PRC1 could not fully re-occupy these loci perhaps due to the presence of SWI/SNF. In agreement with this observation, we also observed an increased level of H3K27ac at the loci where PHC2 could not re-occupy chromatin. Interestingly, while PHC2 and RING1B mostly re-occupied binding sites of wild type cells when PHC2 was re-expressed in the PHC2^KO^ background, H3K27me3 did not recover to wild type levels (**Figure 7F**). The lack of PHC2 re-occupancy and the reduction in H3K27me3 in the PHC2^Re^ line compared to the PHC2^WT^ line correlated with an increase in gene expression levels (**Figure 7G**). This change in gene expression levels might also explain why the gene expression profile of PHC2^Re^ did not recover to PHC2^WT^ levels (**Figure 7D, G**). Taken together, we conclude that PRC1 can re-establish chromatin binding at sites that are not occupied by SWI/SNF but that PRC1 can not reclaim loci that have been occupied by SWI/SNF. This finding in cells is therefore consistent with the *in vitro* reconstitution observations showing that SWI/SNF pre-occupancy on chromatin can restrict PRC1 binding.

## Discussion

Polycomb repressive complexes and the activating chromatin remodeler SWI/SNF are key chromatin regulatory complex families that compete to create repressed and accessible chromatin states, respectively^8–12^. Our findings reveal that the ability of PRC1 to form condensates plays a key role in excluding SWI/SNF from chromatin. The efficiency of this exclusion is determined by the specific complex composition of PRC1. Combinations of PRC1 subunits that enhance condensate formation have an increased ability to block SWI/SNF from chromatin access. In contrast, when SWI/SNF is pre-bound to chromatin, it limits PRC1 binding and the formation of PRC1 condensates. Moreover, we found that SWI/SNF can inhibit PRC1 condensate formation in an ATP-hydrolysis independent manner, indicating a mechanism driven by steric hindrance and competition for chromatin binding rather than active nucleosome remodeling alone. Together, these findings uncover a mutually antagonistic relationship between PRC1 condensates and SWI/SNF complexes. We propose that PRC1 composition and condensate formation play an essential role in restricting SWI/SNF binding to chromatin to maintain transcriptionally silent states (**Figure 7H**). Below, we discuss the details and biological implications of our model in the context of previous findings.

Different PRC1 compositions have strikingly different abilities to restrict SWI/SNF binding to chromatin. Specifically, we show that condensate-promoting PRC1 compositions (e.g. with CBX2) can effectively exclude SWI/SNF from chromatin binding while PRC1 compositions that do not form condensates (e.g. with CBX7) at the same concentrations are quickly evicted by SWI/SNF (**Figure 2, 3**). This observation advances previous work^53–55^ highlighting that the outcome of the PRC1 and SWI/SNF competition is highly context dependent and that not one complex dominates the other. Interestingly, previous work has shown that pluripotent cells, such as mouse embryonic stem cells (mESCs), predominantly express PRC1 subunits that are not associated with condensate formation while more differentiated lineages, such as neural progenitor cells, express higher levels of condensate-promoting PRC1 subunits^17,38,64^. This suggests that as cells progress through differentiation, PRC1 may shift to compositions that are more effective at excluding SWI/SNF to more permanently repress specific genes. We propose that the functional flexibility in PRC1 complex composition is a key mechanism for balancing gene plasticity and stable repression during development.

While this work has shown that PRC1 composition and condensate properties are an important contributor to the antagonism of PRC1 and SWI/SNF, there are likely other mechanisms that add to the opposition. For instance, Sahu et al.^73^ have shown that histone deacetylation contributes to SWI/SNF exclusion from chromatin. The continued maintenance of H3K27me3 by PRC2 and prevention of H3K27ac might thus aid in the prevention of SWI/SNF engagement at Polycomb sites. In addition, functional partitioning of condensates^30,74,75^ might contribute to the separation of PRC1 and SWI/SNF complexes in cells. Patil et al.^33^ have shown that the sequence pattern of the intrinsically disordered region of ARID1A is important to maintain the interaction network of SWI/SNF and is required for correct genomic targeting. While ARID1A condensate formation is mainly driven by tyrosine residues and AQG blocks^33^, PRC1 condensate formation is driven by PHC polymerization and arginine and lysine residues in the CBX subunit^38–41^. These two drivers of condensate formation constitute very different molecular features which might result in partitioning into distinct condensates and thus further contribute to the mutual exclusion of PRC1 and SWI/SNF.

Consistent with the observed antagonism between PRC1 and SWI/SNF in our and in previous *in vitro* reconstitution assays^51,52^, we observed no meaningful colocalization and overlap between the two PRC1 and SWI/SNF complexes in cells. Using super-resolution microscopy we found that PRC1 and SWI/SNF loci are often in close proximity but almost never overlap (**Figure 1**). This is in good agreement with our genomic observations that revealed that only 0.3% of significant PHC2 peaks overlap with significant BAF155 peaks, closely mirroring data by Weber et al.^55^, who observed only 1% overlap between RING1B and significant BAF155 peaks. Moreover, a study investigating the proximal interactome of RING1B in mouse embryonic stem cells^76^ has not found any consensus overlap with SWI/SNF subunits further supporting the lack of overlap of PRC1 and SWI/SNF in cells. Interestingly, when we perturbed PRC1 condensate formation in cells by preventing PHC2 polymerization using the endogenous point mutation L307R or by fully knocking out PHC2, we observed SWI/SNF binding to genes that were previously repressed and bound by canonical PRC1 (**Figure 6, 7**). While SWI/SNF only bound to a few genes that lost canonical PRC1 binding as a result of the point mutation or knockout, most of the genes that were bound by SWI/SNF showed an increase in histone H3 lysine 27 acetylation (H3K27ac) and had an increase in transcription levels. In agreement with this observation, Dobrinić et al. showed increased polymerase II binding and transcription initiation when RING1B was depleted^77^. We propose that PCH2 polymerization and PRC1 condensates are important to restrict SWI/SNF binding to chromatin and to maintain gene repression. Furthermore, we found that PRC1 and SWI/SNF occupy distinct chromatin territories that are mutually exclusive.

While we found that PRC1 can exclude SWI/SNF from accessing chromatin, we also observed the opposite. When SWI/SNF is present first on chromatin, PRC1 can only bind to chromatin to a smaller degree (**Figure 4, 7**). This is in good agreement with Weber el al.^55^ who have shown that only when SWI/SNF is degraded in mESC can PRC1 redistribute to previously SWI/SNF occupied loci but that PRC1 is otherwise kept out. We also observed that condensate formation of PRC1 is abolished if SWI/SNF is present first on chromatin (**Figure 4**). This result aligns with prior studies indicating that PRC1 must extensively coat nucleosomal arrays to drive condensate formation with chromatin^40^. Here, SWI/SNF occupancy on chromatin would simply obstruct PRC1 binding across a given chromatin stretch and thus compromise the multivalent interactions necessary for condensate formation. In agreement with this model, our data showed that PRC1 can re-occupy chromatin at sites with no increase in SWI/SNF levels when PRC1 is re-expressed (**Figure 7**). The observed inhibition of PRC1 binding by SWI/SNF does not rely on changes in nucleosome spacing, as chromatin remodeling per se is not required for PRC1 exclusion (**Figure 5**) suggesting a mechanism driven by competitive binding and steric hindrance rather than active remodeling alone. In agreement with this observation, Moore et al. recently reported that prebinding of the chromatin remodeler ACF or RSC can inhibit nucleosomal array condensate formation in an ATP-independent manner by occluding nucleosome surfaces^44^. This points to a more general mechanism suggesting that the presence of chromatin remodelers can prevent chromatin condensation by steric occlusion of condensate driving factors such as nucleosome surfaces and PRC1 binding. Additionally, Pan et al. previously reported genome-wide catalytic activity dependent and catalytic activity–independent targeting of the SWI/SNF complexes to chromatin^68^. Taken together we propose that SWI/SNF opposes Polycomb repression through both remodeling-dependent and -independent mechanisms to promote chromatin accessibility.

Mutations or altered expression levels of PRC1 and SWI/SNF subunits are linked to a variety of diseases, including developmental disorders and cancers^1–7^. For example, overexpression of the PRC1 subunit CBX2 has been shown to promote cancer progression^78,79^. This elevated CBX2 expression may drive the formation of larger or more stable condensates, leading to excessive gene repression by outcompeting SWI/SNF and disrupting the balance of gene expression. Similarly, mutations in SWI/SNF subunits have been linked to more than 20% of human cancers^80^. One among many effects of SWI/SNF cancer mutations might be that they affect SWI/SNF chromatin binding affinity or impair SWI/SNF condensate formation and thus alter SWI/SNF’s ability to displace PRC1, further tipping the balance between repression and activation. These outcomes are likely subunit- and context-specific, underscoring the importance of condensate dynamics and subunit diversity in understanding the antagonism of Polycomb and SWI/SNF.

## Acknowledgments

We are grateful to all members of the Kingston and Subramanian laboratories (all Massachusetts General Hospital Research Institute and Harvard Medical School, Boston) for critical discussions of this work. We thank Christos G. Tsokos for his advice on generating endogenously tagged cell lines using CRISPR. We are grateful to Ruslan Sadreyev’s lab for access to the sequencing core and Margarete Diaz Cuadros’s lab for providing access to the nucleofector unit (both Massachusetts General Hospital). We thank Luke H. Chao (Massachusetts General Hospital) and Melody Nguyen for advice on the lipid bilayer system. S.N. was supported by the Dennis and Marsha Dammerman Fellowship of the Damon Runyon Cancer Research Foundation (DRG-2418-21) and the NIGMS K99 grant (K99GM155610). The authors gratefully acknowledge funding from NIH grant to R.E.K. (5R35GM131743-07).

## Author Contributions

S.N., R.S., and R.E.K. designed the research; S.N. and S.M. cloned constructs and prepared baculovirus. S.N. expressed and prepared samples; S.N. generated HCT116 cell lines; R.S. performed all REA assays; S.N. collected TIRF, epi, and SIM microscopy data; S.N. performed genomic experiments; S.N. the analyzed microscopy data; P.C.S. and S.N. analyzed genomics data; S.N., R.S., and R.E.K. wrote the manuscript. All authors read and commented on the paper.

## Materials and Methods

### Experimental model and subject details

#### HCT116 cells

(American Type Culture Collection [ATCC]) were cultured in RPMI + GlutaMAX (Thermo Fisher Scientific, 61870-036) supplemented with FBS (Sigma, F0926) to 10% (v/v) concentration and 1% Penicillin Streptomycin (Thermo Fisher Scientific 15140-122). Cells were grown at 37°C with 5% CO_2_ in a humidified sterile incubator.

#### MCF7 cells

(American Type Culture Collection [ATCC]) were cultured in EMEM (Thermo Fisher Scientific, 12440-061) supplemented with FBS (Sigma, F0926) to 10% (v/v) concentration and 1% Penicillin Streptomycin (Thermo Fisher Scientific 15140-122). Cells were grown at 37°C with 5% CO_2_ in a humidified sterile incubator.

#### 4T1 cells

(American Type Culture Collection [ATCC]) were cultured in RPMI + GlutaMAX (Thermo Fisher Scientific, 61870-036) supplemented with FBS (Sigma, F0926) to 10% (v/v) concentration and 1% Penicillin Streptomycin (Thermo Fisher Scientific 15140-122). Cells were grown at 37°C with 5% CO_2_ in a humidified sterile incubator.

#### NIH-3T3 cells

were cultured as previously described^38^. Briefly, NiH-3T3 (American Type Culture Collection [ATCC]) cells were grown in DMEM (Thermo Fisher Scientific, 11995-065) supplemented with 10% fetal bovine serum (FBS) (Sigma, F0926), and 1% Penicillin Streptomycin (Thermo Fisher Scientific 15140-122). Cells were grown at 37°C with 5% CO_2_ in a humidified sterile incubator.

#### J1 Mouse Embryonic Stem Cells

(mESC) (ATCC, SCRC-1010) were cultured on 1% gelatin with a layer of mitotically inactivated mouse embryonic fibroblasts (Millipore; PMEF-N) in ESC medium (DMEM (Thermo Fisher Scientific, 11995-065), 15% (v/v) fetal bovine serum (FBS) (Sigma, F0926), 1× GlutaMAX (Thermo Fisher Scientific, 35050061), 1× penicillin/streptomycin (Thermo Fisher Scientific 15140-122), 1× nonessential amino acids (Thermo Fisher Scientific, 11140050), 0.001% (v/v) 2-Mercaptoethanol, and 10 ng/mL leukemia inhibitory factor (LIF)) (Sigma, ESG1107). ESC media was exchanged daily. Cells were grown at 37°C with 5% CO_2_ in a humidified sterile incubator.

#### HEK293T cells

(American Type Culture Collection [ATCC]) were cultured in IMDM (Thermo Fisher Scientific, 12440-061) supplemented with FBS (Sigma, F0926) to 10% (v/v) concentration. Cells were grown at 37°C with 5% CO_2_ in a humidified sterile incubator.

#### Sf9 cells

(*Spodoptera frugiperda* cell line IPLB-Sf-21-AE) (Expression Systems, 94-001F) were maintained in ESF 921 medium (Expression Systems, 96-001-01) with 1% Penicillin Streptomycin (Thermo Fisher Scientific 15140-122) at 27°C in a shaking incubator.

### Protein expression, purification, and labeling

All PRC1 complexes were expressed in Sf9 cells using the Bac-to-Bac system (Thermo Fisher Scientific, 10359016) as previously described^38,40^. We always expressed all four subunits of canonical PRC1 (RING1B, BMI1, CBX, and PHC) together. For RING1B (mouse), and BMI1 (mouse), we cloned the full length, untagged mouse cDNA into a pBig1a vector^81^ (Addgene, Plasmid #80611). For the CBX subunits (mouse), we cloned the following constructs into a pFastBac1 vector (Thermo Fisher Scientific, 10359016):

Flag-GSAAAGS-mGFP-GSAAAGS-CBX2,

Flag-GSAAAGS-mGFP-GSAAAGS-CBX2-13KRA,

Flag-GSAAAGS-mGFP-GSAAAGS-CBX2-23KRA,

Flag-GSAAAGS-mGFP-GSAAAGS-CBX7. For PHC subunits (mouse), we cloned the following constructs into pFastBac1 vector (Thermo Fisher Scientific, 10359016): PHC2-TEV-HALO and PHC2-L307R-TEV-HALO. The PHC2 subunits have multiple alternative splicing isoforms. We expressed and purified the “short” isoform of PHC2 for all experiments.

For all SWI/SNF complexes we used the following subunits and expressed them simultaneously: BAF250 (ARID1A), Brg1, BAF170, BAF45D, BAF60, BAF57, BAF53, INI1, and ACTB. We cloned the full length, human cDNA of BAF250 into a pFastBac1 vector (Thermo Fisher Scientific, 10359016): ProteinA-GSGSG-TEV-HALO-GSG-BAF250. Moreover, we cloned the full length, untagged human cDNA of Brg1, BAF170, and INI1 into a pBig1a vector^81^ (Addgene, Plasmid #80611). In this vector we also made point mutations to make Brg1 with K785R and D881N. In addition, we cloned the full length, untagged human cDNA of BAF45D, BAF60, BAF57, BAF53, and ACTB into a pBig2ab vector^81^ (Addgene, Plasmid #80616).

We infected Sf9 cells (Expression Systems, 94-001F) with a multiplicity of infection (MOI) of 5 of baculovirus at 1x10^6^ cells per ml and incubated at 27°C for 70 hours in ESF921 (Expression Systems, 96-001-01) 1% Penicillin Streptomycin (Thermo Fisher Scientific 15140-122). Except for virus with Brg1, BAF170, and INI1 for which we used a multiplicity of infection of 50 since Brg1 is very lowly expressed.

For PRC1 and SWI/SNF expression and purification, cells were harvested by centrifugation at 5,000 RCF for 20 min. For PRC1 we typically expressed 4 liters. For SWI/SNF we expressed 12 liters total. The cell pellet of a 1-liter expression was resuspended in 20 ml of PBS at pH 7.4. Afterwards, cells were spun again at 1,000 RCF for 5 min and the supernatant was discarded. We next measured the packed cell volume (PCV) and resuspended the pellet in a hypotonic buffer (10 mM HEPES at pH 7.9, 10 mM KCl, 1.5 mM MgCl_2_, 0.1 mM DTT, 0.2 mM PMSF) to 3x PCV. The cells were incubated on ice for 10 min and lysed with a Dounce homogenizer. Afterwards we centrifuged the lysate at 4,000 RCF for 15 min. Subsequently, we measured the packed nuclear volume (PNV) of the pellet and resuspended the nuclei in a volume of low salt buffer (20 mM HEPES at pH 7.9, 20 mM KCl, 1.5 mM MgCl_2_, 0.2 mM EDTA at pH 8.0, 25% glycerol, 0.1 mM DTT, 0.2 mM PMSF) equal to 1/2 PNV. Then, we added a volume of high salt buffer (20 mM HEPES at pH 7.9, 1200 mM KCl, 1.5 mM MgCl_2_, 0.2 mM EDTA at pH 8.0, 25% glycerol, 0.1 mM DTT, 0.2 mM PMSF) equal to 1/2 PNV and incubated the solution with gentle mixing at 4°C for 30 min. The nuclei were pelleted by centrifugation at 25,000 RCF for 30 minutes. Finally, we flash-froze the supernatant (nuclear extract) and stored it at -80°C.

For PRC1, the nuclear extract was subsequently incubated with MonoRab™ Anti-DYKDDDDK affinity resin (Genscript, L00766) for 2 hours and then washed with BC300 buffer (20 mM HEPES at pH 7.9, 300 mM KCl, 1 mM EDTA, 1 mM MgCl_2_, 10% glycerol, 0.05% IGEPAL CA-630 (Sigma, I8896-100ML), 1 mM DTT, 0.1 mM PMSF, cOmplete EDTA-free protease inhibitor [Roche]). Afterwards the beads were washed with BC600 buffer (20 mM HEPES at pH 7.9, 600 mM KCl, 1 mM EDTA, 1 mM MgCl2, 10% glycerol, 0.05% IGEPAL CA-630, 1 mM DTT, 0.1 mM PMSF, cOmplete EDTA- free protease inhibitor [Roche]) and again with BC300 buffer without IGEPAL CA-630. PRC1 complexes were eluted from the resin using BC300 buffer (without IGEPAL CA-630) containing 0.8 mg/mL Flag peptide.

For SWI/SNF the nuclear extract was subsequently incubated with IgG Sepharose 6 Fast Flow (Cytiva, 17096901) for 4 hours and then washed with BC300 buffer. Afterwards the beads were washed with BC600 buffer and again with BC300 buffer without IGEPAL CA-630. SWI/SNF complexes were eluted by adding TEV protease to 0.05 mg/ml. This mixture was incubated overnight on ice. The next day the elution was collected by harvesting the supernatant.

Subsequently all PRC1 and SWI/SNF complexes were concentrated using Amicon Ultra-4 centrifugal filter units with a MW cutoff of 100 kDa (MilliporeSigma, UFC810024). For SWI/SNF, we labeled HALO-tagged^59^ ARID1A with HALO-TMR dye (Promega, G8251) by incubating the concentrated protein with dye for 1 hour on ice. Afterwards, PRC1 and SWI/SNF complexes were further purified by size exclusion chromatography on a Superose 6 Increase 10/300 GL column (Cytiva, 29-0915-96). Fractions containing full PRC1 or SWI/SNF complexes (**Figure S2C, E**) were concentrated using Amicon Ultra-4 centrifugal filter units with a MW cutoff of 100 kDa (MilliporeSigma, UFC810024) and protein concentration was determined by Bradford assay. Finally, we confirmed the purity of complexes by 4-20% SDS-PAGE (Biorad, #4561096) and Coomassie staining (**Figure S2D, F**). Protein aliquots were flash-frozen and stored at -80°C.

### Preparation of nucleosomal arrays

Recombinant histones were purified following the method described by Dyer et al.^103^ and as previously described^40^. Xenopus histones H2A, H2B, H3, and H4 cloned in a pET28 vector were expressed in E. coli BL21(DE3) cells grown at 37 °C in 2YT medium supplemented with 50 µg/mL kanamycin. Expression was induced with 1 mM IPTG, and cultures were incubated for an additional 3 hours before harvesting by centrifugation at 5,000 RCF for 20 minutes. Cell pellets were resuspended and sonicated in wash buffer (50 mM Tris at pH 7.5, 100 mM NaCl, 1 mM PMSF, 1 mM 2-mercaptoethanol). Inclusion bodies were pelleted by centrifugation at 48,000 RCF for 15 minutes, resuspended in wash buffer containing 1% Triton X-100, and pelleted again at 48,000 RCF for 15 minutes. This wash step was repeated twice. The final pellet was resuspended in SAUDE 200 buffer (7 M urea, 20 mM sodium acetate at pH 5.2, 200 mM NaCl, 5 mM 2-mercaptoethanol, 1 mM EDTA) and centrifuged at 48,000 RCF for 15 minutes. The supernatant was filtered through a 0.22 µm syringe filter (GenClone, 25-243) and loaded onto a 5 mL HiTrap SP HP column (Cytiva, 29-0513-24) pre-equilibrated with SAUDE 200. After washing with 10% SAUDE 1000 (7 M urea, 20 mM sodium acetate at pH 5.2, 1000 mM NaCl, 5 mM 2-mercaptoethanol, 1 mM EDTA), bound proteins were eluted with a linear gradient from 10% to 50% SAUDE 1000. Eluted histone fractions were dialyzed overnight against 5 mM 2-mercaptoethanol and then lyophilized.

The recombinant histone octamer was assembled as previously described^40^. Briefly, purified histones were used as described above, with 4 mg of each histone dissolved in 2 ml unfolding buffer (6 M guanidinium chloride, 20 mM Tris at pH 7.5, 5 mM DTT). The histones were then mixed in equimolar ratios and dialyzed overnight at 4 °C in Slide-A-Lyzer Dialysis Cassettes (7 kDa cutoff; Thermo Fisher Scientific, 66370) against three changes of 1 liter refolding buffer (2 M NaCl, 10 mM Tris at pH 7.5, 1 mM EDTA, 5 mM 2-mercaptoethanol). Precipitated proteins were removed by centrifugation at 48,000 RCF for 15 minutes. The clarified sample was loaded onto a HiLoad 26/600 Superdex 200 gel filtration column (Cytiva, 28-9893-36) pre-equilibrated with refolding buffer. Histone octamer fractions were pooled and stored at 4 °C.

The DNA for nucleosomal arrays were cloned into pUC57 vectors as previously described^40^. We modified the G5E4 plasmid^104,105^ by replacing the ClaI and KpnI restriction sites with BsmBI. The use of BsmBI enabled insertion of non-palindromic sequences, allowing ligation of other dsDNA fragments to the array strands without self-ligation. The modified plasmid was transformed into NEB 5-alpha Competent E. coli (New England Biolabs, C2987H) and cultured in terrific broth (TB) with 100 µg/ml carbenicillin. Cells were harvested by centrifugation at 5,000 RCF for 15 minutes, and plasmid DNA was purified using the Qiagen Plasmid Maxi Kit (Qiagen, 12162) according to the manufacturer’s protocol. To isolate the nucleosomal array DNA from the vector backbone, 8.6 ml of plasmid DNA (10 mg/ml) was mixed with 1.0 ml of 10× CutSmart buffer and 0.2 ml of BsmBI_v2 (NEB, R0739L), then incubated at 55 °C for 48 hours. Next, 0.2 ml of DdeI (NEB, R0175L) was added and the reaction was incubated at 37 °C for an additional 24 hours. Completion of digestion was verified by running 2 µl on a 1% agarose gel; if needed, additional enzyme was added and the reaction was extended by 12 hours. Following digestion, 5 M NaCl was added to a final concentration of 0.5 M. PEG precipitation was then performed by adding PEG-6000 (Sigma, 8074911000) with 0.5 M NaCl to a final PEG concentration of ∼5% (adjusted as needed) and incubating the mixture overnight at 4 °C. The next day, the sample was centrifuged at 29,600 RCF for 20 minutes. The pellet was resuspended in 7 ml TE buffer (10 mM Tris-HCl at pH 8.0, 1 mM EDTA at pH 8.0) and allowed to rehydrate at 4 °C for 4 hours.

To label the DNA with fluorophores or biotin, custom oligonucleotides were ordered from Integrated DNA Technologies (IDT): OligoA-BsmBI_Side1_Biotin (Biotin/TTT TTT TTG GTG TAG GAG GTA GAT GAG); OligoB-BsmBI_Side1_ATTO647N (Phos/ATCG CTC ATC TAC CTC CTA CAC CAA /ATTO647N/); OligoA-BsmBI_Side2_Biotin (Phos/ACTG GGT AGA GTG GTA AGT AGT GAA TTT TTT /Biotin/); OligoB-BsmBI_Side2UN (TTC ACT ACT TAC CAC TCT ACC). For each side, oligos were annealed separately by mixing 9 µl of OligoA (100 µM) with 9 µl of OligoB (100 µM) and 2 µl of 10× DNA Ligase Buffer (NEB, M0202S). The mixtures were heated to 95 °C for 5 minutes, then cooled to 20 °C at 1 °C per minute to form dsDNA fragments with complementary BsmBI overhangs matching the digested array DNA. These labeled dsDNA fragments were ligated to the nucleosomal array strands using T4 DNA ligase (NEB, M0202S) by incubating at 16 °C overnight, with the dsDNA added in 8-fold molar excess relative to the array DNA. Excess dsDNA was removed by performing three rounds of purification with the QIAquick PCR Purification Kit (Qiagen, 28106).

Nucleosomal arrays were assembled as previously described^106^. Briefly, 5 µg of DNA and 5 µg of histone octamer were mixed in high-salt buffer (10 mM Tris-HCl at pH 7.5, 2 M KCl, 1 mM EDTA, 1 mM DTT) and loaded into 7 kDa Slide-A-Lyzer MINI Dialysis Devices (Thermo Fisher Scientific, 69562). Samples were placed in 400 ml of high-salt buffer, and a peristaltic “rabbit” pump was used to gradually exchange the buffer by pumping 1600 ml of low-salt buffer (10 mM Tris-HCl at pH 7.5, 100 mM KCl, 1 mM EDTA, 1 mM DTT) into the beaker while removing buffer at the same rate. Dialysis was carried out overnight at 4 °C, followed by an additional 4-hour dialysis in Storage Buffer (10 mM Tris-HCl at pH 7.5, 10 mM KCl, 1 mM EDTA, 1 mM DTT). The dialyzed nucleosomal arrays were centrifuged at 15,000 RCF for 10 minutes at 4 °C to remove aggregates. Arrays were then digested with BsrFI-v2 (NEB, R0682S, R0161S) for 2 hours at 30 °C and analyzed on a 1% agarose gel to verify nucleosome occupancy on the positioning sequences. Final samples were stored at 4 °C.

### Preparation of lipid bilayers

The lipid bilayers were prepared based on a previously described protocol^40^. Briefly, glass vials (Thermo Fisher Scientific, 14-955-331) and a Hamilton glass syringe (Avanti, 610000-1EA) were cleaned sequentially with Milli-Q water, 70% ethanol, chloroform, 70% ethanol again, and a final rinse with Milli-Q water before air drying. To prepare lipid stocks, 300 mg of 18:1 (Δ9-Cis) PC (DOPC) (Avanti, 850375P-500mg), 25 mg of 18:1 PEG2000 PE (Avanti, 880130P-25mg), and 2.5 mg of 18:1 Biotinyl Cap PE (Avanti, 870273P-25mg) were dissolved in 25 ml of chloroform. Aliquots of 2 ml were prepared, dried under nitrogen, sealed with high-density thread sealant tape, and stored at −20 °C. For use, dried lipid aliquots were resuspended in 2 ml chloroform and could be kept at −20 °C for several weeks. To prepare the working lipid mixture, ∼160 µl of resuspended stock was transferred into a cleaned glass vial using the Hamilton syringe, dried again under nitrogen, and placed in a desiccator overnight. The following day, 1 mL of PBS (pH 7.4) was added to hydrate the dried lipids, yielding a final concentration of 1 mg/ml. The vial was left at room temperature for at least 10 minutes to allow complete resuspension, then sonicated for 5 minutes in a water bath sonicator (Branson 3800). Meanwhile, the lipid extruder (Avanti, 610000-1EA) was assembled, and the syringes were washed with Milli-Q water, 70% ethanol, and Milli-Q water again. The 0.1 µm filter paper (Avanti, 610005-1EA) and support filters (Avanti, 610014-1EA) were pre-wetted with Milli-Q water. After sonication, five freeze–thaw cycles were performed by alternately transferring the vial from liquid nitrogen to ∼40 °C water. The lipid suspension was then extruded by passing it through the extruder 21 times. The final lipid mixture was stored at 4 °C until use. We always freshly prepared bilayers the day of use by starting with the dried lipids and the overnight desiccator storage.

### Flow-cell preparation

The flow-cells were assembled as previously described^40,107^. In brief, double-sided adhesive sheets (Soles2dance, 9474-08×12 — 3M 9474LE 300LSE) were used to cut custom three-channel flow chambers with a paper cutter (Silhouette, Portrait 3). The adhesive cutouts were then used to assemble the flow chambers between standard glass slides (Thermo Fisher Scientific, 12-550-123) and 170 μm thick coverslips (Zeiss, 474030-9000-000). Before assembly, coverslips were cleaned overnight in a 5% v/v solution of Hellmanex III (Sigma, Z805939-1EA) at 50 °C, then rinsed thoroughly with Milli-Q water.

### Preparation of flow-cells with lipid bilayer system

The flow-cells for TIRF imaging were prepared in slightly different ways depending on the experimental design. For most experiments we conducted end-point assays. We typically mixed nucleosomal arrays (5 nM nucleosome concentration final unless otherwise noted) and various concentrations of PRC1 or SWI/SNF in TIRF Buffer (13 mM HEPES at pH 7.9, 100 mM KCl, 1.4 mM MgCl_2_, 0.25 mM EDTA, 0.9 mM ATP, 1% glycerol, 0.04 μg/μl BSA, 0.4 μg/μl kappa casein). We used G5E4 arrays labeled with one ATTO-647N dye (5 nM nucleosome concentration) and two biotins as described above. To all reactions we added 0.01 μg/μl TEV protease. We incubated the 30 µl mixture at 30°C for 30 minutes (unless otherwise noted) in a thermocycler. Afterwards, we added either PRC1 and SWI/SNF to the mixture (depending on which one was added first) and incubated again at 30°C for 30 minutes in a thermocycler (unless otherwise noted). In the meantime, we prepared the flow-cells with lipid bilayers. To this end, 13 μl of lipid bilayer solution was added to the flow chambers, incubated for 2 minutes, and washed three times with 13 μl of TIRF buffer. Next, 10 μl of streptavidin (Vector, SA-5000) was added and incubated for 2 minutes, followed by two washes with 13 μl of TIRF buffer. Afterwards, we added 13 μl of the array - PRC1 - SWI/SNF sample and incubated for 4 minutes. We then added an additional 13 μl of the array - PRC1 - SWI/SNF sample and incubated for 4 minutes and then started imaging.

For live imaging of condensate formation and SWI/SNF competition (**Figure 2B, C**), we simultaneously mixed nucleosomal arrays (5 nM nucleosome concentration), 5 nM SWI/SNF, and 40 nM of PRC1 in TIRF Buffer. To all reactions we added 0.01 μg/μl TEV protease. This mixture was added to the imaging chamber, which was prepared before mixing the samples) and imaged immediately (∼1 minute delay).

For all experiments, it is important to note that the ratio of ATP, magnesium, and EDTA is critical. Even slight variations in these concentrations can lead to different outcomes (for example, excessive magnesium can cause the arrays to condense on their own).

### Preparation of flow-cells for epi-fluorescence microscopy (droplet assays)

We typically prepared reactions by mixing nucleosomal arrays (30 nM nucleosome final concentration, unless otherwise noted) with varying concentrations of PRC1 and/or SWI/SNF in Droplet Assay Buffer (13 mM HEPES at pH 7.9, 120 mM KCl, 1.4 mM MgCl_2_, 0.25 mM EDTA, 0.9 mM ATP, 1% glycerol, 0.04 μg/μl BSA, 0.4 μg/μl kappa casein). All experiments used G5E4 arrays labeled with ATTO-647N dye. Each reaction also contained 0.01 μg/μl TEV protease. The 30 μl mixtures were incubated at 30 °C for 30 minutes in a thermocycler. Next, PRC1 or SWI/SNF (depending on the experimental order) was added and incubated for an additional 30 minutes at 30 °C. Meanwhile, flow-cells were prepared by washing twice with Droplet Assay Buffer. After the final incubation, 13 μl of the sample was loaded into the flow-cell and incubated for 4 minutes. An additional 13 μl of the array–PRC1–SWI/SNF mixture was then added, followed by another 4-minute incubation before imaging commenced.

### Restriction Enzyme Accessibility (REA) assay

The REA assay was performed similarly to how it has been previously described^45^. Briefly, we first mixed nucleosomal arrays (1.7 nM nucleosome concentration) and various concentrations of PRC1 in REA Buffer (13 mM HEPES at pH 7.9, 100 mM KCl, 1.4 mM MgCl_2_, 0.25 mM EDTA, 0.9 mM ATP, 1% glycerol, 0.04 μg/μl BSA). To all reactions we added 0.01 μg/μl TEV protease. This reaction was incubated at 30°C for 30 minutes in a thermocycler. Then we added 5 nM SWI/SNF (unless otherwise noted) to the reactions and added the restriction enzyme HhaI to 8 units per reaction (unless otherwise noted). Then, this reaction was incubated at 30°C for 60 minutes in a thermocycler. Afterwards, the reaction was quenched by adding 10 μl of STOP buffer (10 mM Tris at pH 7.7, 70 mM EDTA, 1% SDS, 0.1% orange G) to the 30 μl reaction. This mixture was incubated at 55°C for 10 minutes in a thermocycler. Subsequently, the samples were run a 1% agarose gel and scanned with Typhoon biomolecular imager to visualize dye-labeled DNA. Intensities of bands were quantified using the Gel Analyzer in Fiji (light microscopy data)^84^.

### Cell culture and CRISPR generation of cell lines

HCT116 cells (American Type Culture Collection [ATCC]) were cultured in RPMI + GlutaMAX (Thermo Fisher Scientific, 61870-036) supplemented with FBS (Sigma, F0926) to 10% (v/v) concentration and 1% Penicillin Streptomycin (Thermo Fisher Scientific 15140-122). Cells were grown at 37°C with 5% CO_2_ in a humidified sterile incubator.

To endogenously-tag PHC2 and ARID1A we used the CRISPR-Cas9 based PITCh vector system^82,108^ and inserted mEOS3.2^60^ at the N-terminus of ARID1A (puromycin resistance; homology arms of 20 bp). Similarly, we tagged PHC2 with a HALO-tag^59^ at its N-terminus (blasticidin resistance), specifically labeling the long PHC2 isoform (homology arms of 20 bp). We also created the endogenous mutation L307R for PHC2 in HCT116 cells. For the endogenous PHC2 L307R mutation Alt-R HDR Donor template was ordered from IDT (homology arms of 100 bp). To create the PHC2 knockout cell line in HCT116 cells, we used a guide RNA targeting the start codon of the short isoform which also leads to an internal cut of the long PHC2 isoform.

For all cell lines, the CRISPR guide RNA (gRNA) was designed by identifying the nearest PAM sequence (5′-NGG) to the desired insertion site. The 20 bp sequence immediately upstream of this PAM was selected as the guide sequence, following the method described by Ran et al.^109^. Alt-R gRNAs and the universal tracrRNA were obtained from IDT.

To assemble the CRISPR-RNP complex, 10 µl of each 200 µM gRNA and 200 µM tracrRNA were annealed by gradual cooling from 95°C. For PHC2 L307R and PHC2 knockout, 5 µl of the hybridized gRNA (∼250 pmol) was combined with ∼155 pmol of Cas9 protein and incubated at room temperature for 30 minutes. For endogenous tagging of PHC2 and ARID1A, 2.5 µl of the hybridized gRNA (∼125 pmol) was combined with ∼75 pmol of Cas9 protein. At the same time 2.5 µl of PITCh gRNA (∼125 pmol) was combined with ∼75 pmol of Cas9 protein. Both mixtures were incubated at room temperature for 30 minutes. Prior to transfection, 150 ng of linear hygromycin resistance marker (only for PHC2 L307R and PHC2 knockout) and ∼1 µg of donor plasmid or linear donor template were added to the mixture.

For each transfection, 1 × 10^6^ cells were resuspended in 100 µl of nucleofection solution (Lonza Bioscience, V4XC-2024) and mixed with the CRISPR-RNP solution. Immediately after, cells were nucleofected using SF Cell Line 4D-Nucleofector™ X Kit L (Lonza Bioscience, V4XC-2024) in a Lonza 4D-Nucleofector™ unit. Afterwards cells, were plated in RPMI + GlutaMAX (Thermo Fisher Scientific, 61870-036) supplemented with FBS (Sigma, F0926) to 10% (v/v) concentration and 1% Penicillin Streptomycin (Thermo Fisher Scientific 15140-122). After 1 day, selection was started with either 0.5 µg/ml puromycin, 10 µg/ml blasticidin, or 300 µg/ml hygromycin (all Invivogen) for seven days. Individual clones were picked, screened via PCR, and confirmed by Sanger sequencing and western blot.

Lentivirus for the re-expression of PHC2 was made by using HEK293T cells (American Type Culture Collection, ATCC) in IMDM (Thermo Fisher Scientific, 12440-061) supplemented with FBS (Sigma, F0926) to 10% (v/v) concentration. Cells were grown at 37°C with 5% CO_2_ in a humidified sterile incubator. To express the short isoform of human PHC2, the cDNA was inserted into a pTRIPZ vector (Dharmacon). This construct was co-transfected into HEK293T cells along with second-generation lentiviral packaging plasmids—pCMV-dR8.91 (carrying gag, pol, and rev genes) and pMD2.G (encoding the VSV-G envelope protein)—using the TransIT-Lenti transfection reagent (Mirus, MIR 6650). Viral supernatant was collected 48 hours post-transfection, filtered through a 0.45 µm membrane, and used to transduce HCT116 cells with the PHC2 knockout background. Forty-eight hours after infection, cells were selected with 1 µg/ml puromycin. Expression of PHC2 was confirmed by western blot and a doxycycline concentration was selected that results in expression levels similar to those of PHC2 in wild type HCT116 cells. For all RNA sequencing, CUT&RUN, immunoblot, and immunofluorescence microscopy experiments, PHC2 expression was induced by adding doxycycline (1 µg/ml) for 48 hours.

Cells for live-cell imaging and immunofluorescence microscopy were plated at 0.2 × 10^5^ cells per ml in Poly-L-Lysine coated 8-well µ-Slide chambers (Ibidi, 80804). For Structured Illumination Microscopy cells were plated at 0.2 × 10^5^ cells per ml in μ-Slide 8 well high Glass chamber (Ibidi, 80827) that was coated with poly-D-Lysine before. To perform live cell imaging with HALO-JFX549^110^, the HALO dye was diluted to a final concentration of 200 nM in FluoroBrite (Thermo Fisher Scientific, A1896701) imaging media and incubated for 15 minutes. Afterwards, the cells were washed three times with Fluorobrite medium to remove unbound dye. Then the cells were incubated at 37°C for another 15 minutes and the media was replaced with FluoroBrite imaging media supplemented with 20 mM HEPES at pH 7.4. For immunofluorescence microscopy, we fixed the cells after 24 hours (details below).

### TIRF and epi-fluorescence microscopy data collection

The TIRF and epi-fluorescence data collections were conducted using a Nikon Ti-E inverted microscope equipped with a Ti-ND6-PFS perfect focus system and an APO TIRF 100x oil/1.49 DIC objective (Nikon). The microscope was also equipped with a Nikon-encoded x-y motorized stage and a piezo z-stage, an EMCCD camera (Andor iXon Ultra, DU-897U-CSO-#BV), and 488 nm, 561 nm and 640 nm laser. All imaging was performed with subsequent exposures of 102 msec, using the CCD mode (i.e. no EM gain) with 3 MHz and 16-bit. For all imaging we used cubes specifically for each laser line to minimize crosstalk (Chroma, TRF49904 (488 nm), TRF49909 (561 nm), TRF49914 (640 nm)).

### Confocal microscopy and Structure Illumination Microscopy of HCT116 cells

Images of HCT116 cells were acquired on a Nikon A1R laser-scanning confocal inverted microscope equipped with a 100× oil immersion objective. We collected data with a scanning speed of 1/16 frames /second over an area of 1024x1024 pixels and with 8x averaging. We selected “channel series” to avoid crosstalk and set the pinhole to 0.8. For all live-cell imaging the 488 nm laser was set to HV=120, Offset=0, and Power=5.00. The 561 nm laser was set to HV=120, Offset=0, and Power=5.00. For immunofluorescence microscopy, the data was collected at room temperature with 4x averaging and a zoom of 3.0. We set the 405 nm to HV=100, Offset=0, and Power=1.00, the 488 nm laser to HV=100, Offset=0, and Power=1.00, and the 561 nm laser to HV=80, Offset=0, and Power=5.00.

For Structured Illumination Microscopy we used the Nikon N-SIM-E Super Resolution Microscope system with a 100x oil immersion objective. Before each imaging session the light paths were aligned. We used the 12 bit sensitive camera setting with 20 msec exposure for 488 nm and 640 nm (both 100% laser power). We set the ROI 1024 × 1024 and used 0.12 µm steps over the range of 5 µm for the z-stacks. For image reconstruction, Nikon Elements was used by adjusting “Illumination Modulation Contrast”, “High Resolution Noise Suppression”, and “Out of Focus Blur Suppression” ensuring that no artifacts were created.

### Immunofluorescence microscopy of HCT116 cells

HCT116 cells were plated as described above. After 24 hours, cells were rinsed three times with PBS (pH 7.4), fixed in 4% paraformaldehyde for 10 minutes at room temperature, and washed three times with PBS for 10 minutes each. Cells were then permeabilized with 0.1% Triton X-100 in PBS for 10 minutes at room temperature, followed by three additional PBS washes (10 minutes each). Blocking was performed for 20 minutes at room temperature in a solution of 1% BSA and 0.1% Tween-20 in PBS. Cells were then incubated overnight at 4 °C with the primary antibody diluted 1:400 in blocking solution. The next day, cells were washed three times with blocking solution for 10 minutes each and incubated with the secondary antibody (1:1,000 in blocking solution) for 2 hours at room temperature in the dark. Afterward, cells were washed once with blocking solution, stained with Hoechst dye (1:10,000) for 10 minutes, and washed three final times with blocking solution for 10 minutes each.

### RNAseq sample and library preparation

For all RNA sequencing experiments we grew HCT116 cells in 6-well plates. The day before harvesting the cells, we plated 2.5×10^5^ cells per well. Cells were harvested by adding 0.5 ml of TRI Reagent (Molecular Research Center, TR118) and transferred to a 1.5 ml tube. Then 0.1 ml of chloroform was added to the tube and vigorously mixed. Afterwards, the sample was incubated at room temperature for 3 minutes and centrifuged at 16,000 RCF for 15 minutes at 4°C. We then carefully transferred the upper aqueous phase to a new tube and added 1 volume of 100% ethanol to 1 volume of sample (1:1 ratio). Then, the RNA was purified using the RNA Clean & Concentrator-5 kit (Zymo Research, R1016) following the manufacturer’s manual. To remove the DNA we performed the “In-column DNase treatment” and used 5 units of TURBO DNAase (Thermo Fisher Scientific, AM2238) as described in the appendix of the manufacturer manual. The quality and concentration was accessed by nanodrop and Agilent TapeStation.

For the experiments comparing HCT116 cells with PHC2 wild type and PHC2 L307R (**Figure 6**), the RNA library was prepared using 250 ng of RNA and the Zymo-Seq RiboFree Total RNA Library Kit (Zymo Research, R3000) following the manufacturer’s manual exactly. For the final library amplification we ran 12 PCR cycles. Afterwards library quality was assessed using the Agilent TapeStation. The library concentrations were measured using the NEBNext Library Quant Kit for Illumina (New England Biolabs, E7630L) following the manufacturer’s manual exactly.

For the experiments comparing HCT116 cells with PHC2 wild type, PHC2 knockout, and the PHC2 re-expression (**Figure 7**), the RNA library was prepared using 800 ng of RNA and the NEBnext PolyA mRNA magnetic isolation module (New England Biolabs; E7490L) followed by the NEBnext ultraii directional RNA library prep (New England Biolabs, E7760L). For both, we followed the manufacturer’s manual exactly. We diluted the adapter 5-fold and for the final library amplification we ran 12 PCR cycles. Afterwards library quality was assessed using the Agilent TapeStation. The library concentrations were measured using the NEBNext Library Quant Kit for Illumina (New England Biolabs, E7630L) following the manufacturer’s manual exactly.

### CUT&RUN sample and library preparation

We followed the CUT&RUN protocol from Skene et al.^61^ with some minor modifications. First, Concanavalin A (Polysciences, 86057-10) beads were prepared using 20 µl per sample and washing the beads three times with 1 ml of Bead Activation Buffer (20 mM HEPES with pH 7.9, 10 mM KCl, 1 mM CaCl_2_ 1 mM MnCl_2_), incubating for ∼15 minutes on a rotator each time. After washing, tubes were placed on a magnetic stand, supernatant was removed, and beads were resuspended in 200 µl Bead Activation Buffer.

For each CUT&RUN sample, 1×10^6^ cells were harvested by trypsinization, quenched, and counted. Then cells were spun down at 500 RCF for 5 minutes and washed with 10 ml of PBS at pH 7.4. We resuspended cells in 0.5% paraformaldehyde in PBS at pH 7.4. Cells were fixed for 5 minutes, then quenched with 1:1 volume of 1 M Tris at pH 8.0 for 10 minutes at room temperature. Afterwards, we washed cells twice with 5 ml PBS pH 7.4 in 15 ml Falcon tubes by spinning at 500 RCF for 3 minutes. Cells were resuspended in 100 µl of PBS at pH 7.4 per library and we added 100 µl Wash Buffer (20 mM HEPES pH 7.5, 150 mM NaCl, 2 mM EDTA pH 8.0, 0.5% BSA, 0.1% Triton, Protease Inhibitor Tablet Roche cOmplete EDTA-free) and 40 µl of washed ConA beads. This mixture was incubated for 10 minutes on a rotator to bind cells to beads. Then, samples were spun at 50 RCF for 1 minute, moved to a magnetic stand, and cleared for 1 minute. Afterwards, the supernatant was removed and beads were resuspended in 100 µl Wash Buffer and transferred to PCR strip tubes. We then prepared the corresponding antibody solutions (all antibodies were used at 1:400) in 150 µl Wash Buffer. PCR tubes with beads were moved to a magnetic stand, incubated for 1 minute, and then the supernatant was removed. We then added the antibody solution and resuspended the beads. Finally, the tubes were placed on a nutator overnight at 4°C.

The next day, the samples were washed twice with 100 µl Wash Buffer. After removing the Wash Buffer, we added 50 µl of proteinA MNase at 700 ng/ml final in Wash Buffer and incubated for 1 hour at room temperature. Then, the beads were washed three times with 100 µl Wash Buffer. Afterwards, the samples were chilled on ice. We then added 100 µl of ice cold Wash Buffer with 10 mM CaCl_2_ to each sample and digested for 1 hour at 4°C. During the incubation, we prepared RNase-containing 2× STOP buffer by mixing 4 µl of 100 mg/ml RNase A with 3996 µl 2× STOP buffer (340 mM NaCl, 20 mM EDTA pH 8.0, 4 mM EGTA pH 8.0). After the 1-hour incubation, we added 100 µl of STOP/RNase buffer per sample and incubated at 37°C for 15 minutes. Then, samples were placed on the magnet, allowed to clear for 1 minute, and the supernatant was transferred to new tubes. We then added 2 µl of 10% SDS and 2 µl of 19 mg/mL proteinase K to each tube and mixed gently. Reverse crosslinking was performed at 50°C overnight.

The next day, we performed the DNA Clean-Up using Ampure XP beads (Beckman Coulter, A63881) at 0.9x beads to sample ratio (e.g. 180 µl of beads with 200 µl of DNA sample). Beads and samples were mixed and incubated at room temperature for 10 minutes. Afterwards, we moved the tubes to the magnet, discarded the supernatant, and washed with 200 µl of freshly prepared 90% ethanol without resuspending. This wash was repeated and then the beads were allowed to air dry just until no visible liquid remained. We then removed the tubes from the magnet, added 52 µl of 0.1× TE, resuspended beads by pipetting, and incubated for 2 minutes at room temperature. Then, the tubes were returned to the magnet and we transferred 50 µl of supernatant to new PCR tubes.

Afterwards, we continued with library preparation using the NEBNext Ultra II DNA Library Prep Kit (New England Biolabs, E7645), following manufacturer’s instructions with the following modifications. At every step we used only half of the recommended volume. For the adapter ligation we used an adapter ligation dilution of 1:50. After the adapter ligation, the sample was cleaned up using Ampure XP beads at 1.2x beads to sample ratio. For the library amplification, we ran 16 cycles. The amplified library was cleaned with Ampure XP beads at 1.2x beads to sample ratio. Finally, the library quality was assessed using the Agilent TapeStation. The library concentrations were measured using the NEBNext Library Quant Kit for Illumina (New England Biolabs, E7630L) following the manufacturer’s manual exactly.

### Immunoblots

Cells were cultured as described in the **Experimental Model and Subject Details**, except for J1 Mouse Embryonic Stem Cells (mESC) (ATCC, SCRC-1010), which were maintained on 1% gelatin-coated plates layered with mitotically inactivated mouse embryonic fibroblasts (Millipore; PMEF-N) in ESC medium. The ESC medium consisted of DMEM (Thermo Fisher Scientific, 11995-065), 15% (v/v) fetal bovine serum (FBS) (Sigma, F0926), 1× GlutaMAX (Thermo Fisher Scientific, 35050061), 1× penicillin/streptomycin (Thermo Fisher Scientific, 15140-122), 1× nonessential amino acids (Thermo Fisher Scientific, 11140050), 0.001% (v/v) 2-mercaptoethanol, and 10 ng/mL leukemia inhibitory factor (LIF) (Sigma, ESG1107). ESC medium was replaced daily. Neural progenitors (NPCs) were generated following a previously described protocol^64^. Initially, mESCs were passaged twice to remove feeder cells (de-MEFed). Then, 4 × 10^6^ cells were seeded into low-adherence 10 cm dishes containing 15 mL of CA medium composed of DMEM supplemented with 10% FBS, 2 mM L-glutamine (Thermo Fisher Scientific, 25030081), 1× nonessential amino acids, 1× penicillin/streptomycin, and 2-mercaptoethanol. The medium was refreshed after 2 days. Four days after the initial transfer to CA medium, the medium was replaced with CA medium supplemented with retinoic acid at a final concentration of 5 μM (Sigma, R2625-50MG). After an additional two days, the medium was replaced again with CA medium containing 5 μM retinoic acid. Cells were harvested on day 8 following the first media change.

All cells were harvested 24 hours after plating by trypsinization, followed by centrifugation at 600 RCF for 5 minutes. Pellets were washed with PBS (pH 7.4) and centrifuged again at 600 RCF for 5 minutes. Subsequently, pellets were resuspended in 100 μl RIPA buffer (Thermo Fisher Scientific, 89900), and protein concentrations were measured using a BCA Assay (Thermo Fisher Scientific, 23225). Protein lysates were separated on 4–20% SDS-PAGE gels (Biorad, 5671095) and transferred to 0.45 μm PVDF membranes (Millipore, IPFL00010) using a Mini Trans-Blot Cell (Biorad, 1703930) with cold transfer buffer (25 mM TRIS Base, 192 mM glycine, 20% methanol) at 350 mA for 75 minutes at 4°C. Following transfer, membranes were blocked in 2% milk in TBST (20 mM TRIS at pH 7.8, 150 mM NaCl, 0.1% Tween-20) for 1 hour at room temperature. Membranes were then incubated overnight at 4°C in primary antibodies diluted 1:1,000 in 2% milk/TBST. After three washes with TBST at room temperature for 10 minutes each, membranes were incubated for 2 hours at room temperature with secondary antibodies diluted 1:10,000 in 2% milk/TBST. The secondary antibodies used included Goat Anti-Mouse IgG IRDye 680 Conjugated (LI-COR Biosciences, 926-32220), Goat Anti-Mouse IgG DyLight™ 800 (Thermo Fisher Scientific, SA5-10176), Goat Anti-Rabbit IgG DyLight™ 680 (Thermo Fisher Scientific, 35569), and Goat Anti-Rabbit IgG Alexa Fluor™ Plus 800 (Thermo Fisher Scientific, A32735). After three final washes with TBST for 10 minutes each, membranes were scanned using an Odyssey CLx Imager (LI-COR).

### TIRF and epi-fluorescence microscopy data analysis

The raw microscopy data was analyzed as previously described^40^. Briefly, all files were analyzed using Fiji (for light microscopy data) and Python (version 3.9.6, Python Software Foundation). To extract spot intensity values from all images, we developed a custom Fiji macro. First, images were opened with Bio-Formats and converted to 8-bit. Background subtraction was performed, followed by applying an automatic intermodes dark threshold to create a mask for all spots. The Watershed algorithm was then applied to separate adjacent spots that appeared merged. Using this mask, raw intensity values, as well as area and perimeter measurements for each spot, were collected and exported as.txt files. These measurements allowed us to calculate roundness for each spot. Additionally, background intensity was determined by measuring the intensity of the entire image excluding the masked spots, and these values were also saved as.txt files. The exported data were further processed with a custom Python script, where the background intensity was divided into 36 equal regions to estimate local background levels more accurately. Background intensity was then subtracted from each spot’s measured intensity. These background-corrected values were used to generate the graphs presented in this manuscript.

### Analysis of cell imaging data

All data was analysed using Nikon Elements and Fiji (light microscopy data). We used the “Spot intensity” plug-in (https://imagej.net/plugins/spot-intensity-analysis) to count the number of spots per cell (**Figure 6D, 7C**) with the following settings: Time interval of 1.0 s, Electrons per ADU of 1.0, Check first n frames of 1, Spot radius of 6, Noise tolerance of 80, and Background estimation Median. Subsequently, the data was further analyzed and plotted using a custom python script. We also used Fiji (light microscopy data) to perform the line scans of various intensities and plotted the raw data using Numbers (Apple Inc). To determine the colocalization of ARID1A and PHC2 (Pearson Correlation Coefficient) in the nucleus (**Figure 1C**), we drew a rectangular ROI around each nucleus and then used the Colocalization Plugin in Nikon Elements. The data was exported and plotted using a custom Python script. For the image reconstruction from the Structured Illumination Microscopy data, Nikon Elements was used by adjusting “Illumination Modulation Contrast”, “High Resolution Noise Suppression”, and “Out of Focus Blur Suppression” ensuring that no artifacts are created.

### Analysis of RNAseq data

The raw reads were initially trimmed using Trimmomatic with the empirically determined parameters “HEADCROP:0 ILLUMINACLIP:adapters.fa:2:22:9:2:True LEADING:3 TRAILING:3 SLIDINGWINDOW:4:15 MINLEN:20”^86^. Quality control of the sequencing data was performed with FastQC and MultiQC^87^.

Subsequently, the trimmed reads were aligned to the ribosomal DNA references, retrieved from NCBI with bowtie2 using the -N 1 option to allow single mismatches in the alignment^88^. Reads identified as non-ribosomal were subjected to splicing-aware alignment with the STAR aligner using the settings “--bamRemoveDuplicatesType UniqueIdenticalNotMulti --outMultimapperOrder Random”. For this step, we employed the hg38 genome and RefSeq annotations for introns and exons^95,96^.

Unaligned reads were removed for further processing and a subset of the unaligned reads was used with BLAST to confirm the absence of any contamination. Picard’s markDuplicate function was used on the aligned read set to identify duplicate reads (https://broadinstitute.github.io/picard/).

Using the Bedtools’ genomecov function, we generated bedgraph files from BAM files with single-base-pair resolution^92^. These files were normalized to 1x genome coverage and converted to bigwigs. In parallel to this, BAM files were processed by featureCounts to determine gene-and-exon-wise coverage in all RNA-Seq samples^94^.

The count table was further processed in R using RStudio (R Core Team, 2023; RStudio Team, 2020). Differential gene expression analysis was conducted with egdeR^100^ and fgsea^99^ was used with gene sets from MSigDB^102^ for gene set enrichment analysis. Gene and exon counts were normalized using the edgeR package’s TMM method^100^ and MDS plots were generated via the limma package^98^.

### Analysis of CUT&RUN data

Raw CUT&RUN reads were trimmed with Trimmomatic^86^ and empirically determined parameters “HEADCROP:5 ILLUMINACLIP:adapters.fa:2:22:9:2:True LEADING:3 TRAILING:3 SLIDINGWINDOW:4:15 MINLEN:20”. Quality control of the CUT&RUN sequencing experiment was performed with FastQC and MultiQC^87^. Mitochondrial reads were removed by alignment to the mitochondrial DNA sequence reference from the hg38 genome using Bowtie2.

The remaining non-mitochondrial reads were further aligned to the human genome hg38 using Bowtie 2, allowing for single-base mismatches (parameter -N 1)^88^. Poor alignments in the produced BAM alignment file, as determined by score fields AS, YS and MAPQ, multi-aligners, as determined by fields AS and XS, large fragments (TLEN>3500) and flagged singletons were subsequently filtered out with samtools^90^.

Replicate marking and removal were accomplished using Picard’s MarkDuplicates (https://broadinstitute.github.io/picard/). Sample- and antibody-specific CUT&RUN peaks were called with macs3^91^, employing multiple replicates for each experiment and using respective IgG CUT&RUNs as controls with a q-value of 0.01 (-q 0.01).

For normalizing CUT&RUN tracks, a “background-interval-set” was generated using the complement of a union peak set derived from all samples and antibodies, extending each peak interval by 10 kbp. Bedgraph files, prepared for each sample with BedTools^92^ genomecov, were normalized to a standard 1x coverage on the background intervals.

Binding profiles and binding heatmaps shown in the figures were generated with deeptools plotHeatmap function^93^.

### Integration of RNAseq and CUT&RUN experiments

The integration of CUT&RUN data with RNA-Seq data required merging peak sets on an antibody-specific basis. Using featureCounts^94^, we counted the number of fragments from the CUT&RUN samples that covered each antibody-specific consensus-peak. We then moved forward with the consensus peak set and peak-based featureCount tables in an R environment. The findOverlaps function from the GenomicRanges package^101^ was utilized to produce pairs of CUT&RUN peaks and promoters. In this process, exon-specific RNA-seq read counts (RPKM-normalized) allowed identifying active transcription start sites (TSSs) for each gene. These TSSs were extended by 1.5 kbp, both upstream and downstream, to encompass promoter regions. Like differential expression analysis, differential CUT&RUN analysis also leveraged the afore-mentioned edgeR package.

### Figure preparation

All figures and graphs were prepared by either using Fiji (light microscopy data), Affinity designer (version 1.6.1, Serif (Europe) Ltd), Python (version 3.9.6, Python Software Foundation) R (R Core Team), or the Integrative genomics viewer^85^.

### Quantification and statistical analysis

The exact values of the number of samples analyzed and the number of repeats per experiment (technical or biological replicates) for each experiment can be found in the figure legends. Error bar representation is indicated in the figure legends.

## References

1. Flavahan, W.A., Gaskell, E., and Bernstein, B.E. (2017). Epigenetic plasticity and the hallmarks of cancer. Science 357. 10.1126/science.aal2380.

2. Alfert, A., Moreno, N., and Kerl, K. (2019). The BAF complex in development and disease. Epigenetics Chromatin 12, 19. 10.1186/s13072-019-0264-y.

3. Valencia, A.M., and Kadoch, C. (2019). Chromatin regulatory mechanisms and therapeutic opportunities in cancer. Nat. Cell Biol. 21, 152–161. 10.1038/s41556-018-0258-1.

4. Mittal, P., and Roberts, C.W.M. (2020). The SWI/SNF complex in cancer — biology, biomarkers and therapy. Nat. Rev. Clin. Oncol., 1–14. 10.1038/s41571-020-0357-3.

5. Piunti, A., and Shilatifard, A. (2021). The roles of Polycomb repressive complexes in mammalian development and cancer. Nat. Rev. Mol. Cell Biol. 22, 326–345. 10.1038/s41580-021-00341-1.

6. Parreno, V., Martinez, A.-M., and Cavalli, G. (2022). Mechanisms of Polycomb group protein function in cancer. Cell Res., 1–23. 10.1038/s41422-021-00606-6.

7. Gourisankar, S., Krokhotin, A., Wenderski, W., and Crabtree, G.R. (2024). Context-specific functions of chromatin remodellers in development and disease. Nat. Rev. Genet. 25, 340–361. 10.1038/s41576-023-00666-x.

8. Kingston, R.E., and Tamkun, J.W. (2014). Transcriptional Regulation by Trithorax-Group Proteins. Cold Spring Harb. Perspect. Biol. 6, a019349–a019349. 10.1101/cshperspect.a019349.

9. Kadoch, C., Copeland, R.A., and Keilhack, H. (2016). PRC2 and SWI/SNF Chromatin Remodeling Complexes in Health and Disease. Biochemistry 55, 1600–1614. 10.1021/acs.biochem.5b01191.

10. Schuettengruber, B., Bourbon, H.-M., Croce, L.D., and Cavalli, G. (2017). Genome Regulation by Polycomb and Trithorax: 70 Years and Counting. Cell 171, 34–57. 10.1016/j.cell.2017.08.002.

11. Bracken, A.P., Brien, G.L., and Verrijzer, C.P. (2019). Dangerous liaisons: interplay between SWI/SNF, NuRD, and Polycomb in chromatin regulation and cancer. Genes Dev. 10.1101/gad.326066.119.

12. Kuroda, M.I., Kang, H., De, S., and Kassis, J.A. (2020). Dynamic Competition of Polycomb and Trithorax in Transcriptional Programming. Annu. Rev. Biochem. 89, annurev-biochem-120219-103641. 10.1146/annurev-biochem-120219-103641.

13. Kennison, J.A., and Tamkun, J.W. (1988). Dosage-dependent modifiers of polycomb and antennapedia mutations in Drosophila. Proc. Natl. Acad. Sci. 85, 8136–8140. 10.1073/pnas.85.21.8136.

14. Tamkun, J.W., Deuring, R., Scott, M.P., Kissinger, M., Pattatucci, A.M., Kaufman, T.C., and Kennison, J.A. (1992). brahma: A regulator of Drosophila homeotic genes structurally related to the yeast transcriptional activator SNF2SWI2<math><mtext>SNF</mtext><mtext>2</mtext> <mtext>SWI</mtext><mtext>2</mtext></math>. Cell 68, 561–572. 10.1016/0092-8674(92)90191-E.

15. Yu, B.D., Hess, J.L., Horning, S.E., Brown, G.A.J., and Korsmeyer, S.J. (1995). Altered Hox expression and segmental identity in Mll-mutant mice. Nature 378, 505–508. 10.1038/378505a0.

16. Blackledge, N.P., and Klose, R.J. (2021). The molecular principles of gene regulation by Polycomb repressive complexes. Nat. Rev. Mol. Cell Biol. 22, 815–833. 10.1038/s41580-021-00398-y.

17. Kim, J.J., and Kingston, R.E. (2022). Context-specific Polycomb mechanisms in development. Nat. Rev. Genet. 23, 680–695. 10.1038/s41576-022-00499-0.

18. Centore, R.C., Sandoval, G.J., Soares, L.M.M., Kadoch, C., and Chan, H.M. (2020). Mammalian SWI/SNF Chromatin Remodeling Complexes: Emerging Mechanisms and Therapeutic Strategies. Trends Genet. 36, 936–950. 10.1016/j.tig.2020.07.011.

19. Bieluszewski, T., Prakash, S., Roulé, T., and Wagner, D. (2023). The Role and Activity of SWI/SNF Chromatin Remodelers. Annu. Rev. Plant Biol. 74, 139–163. 10.1146/annurev-arplant-102820-093218.

20. Mashtalir, N., D’Avino, A.R., Michel, B.C., Luo, J., Pan, J., Otto, J.E., Zullow, H.J., McKenzie, Z.M., Kubiak, R.L., St. Pierre, R., et al. (2018). Modular Organization and Assembly of SWI/SNF Family Chromatin Remodeling Complexes. Cell 175, 1272-1288.e20. 10.1016/j.cell.2018.09.032.

21. Zhou, C.Y., Johnson, S.L., Gamarra, N.I., and Narlikar, G.J. (2016). Mechanisms of ATP-Dependent Chromatin Remodeling Motors. Annu. Rev. Biophys. 45, 153–181. 10.1146/annurev-biophys-051013-022819.

22. Clapier, C.R., Iwasa, J., Cairns, B.R., and Peterson, C.L. (2017). Mechanisms of action and regulation of ATP-dependent chromatin-remodelling complexes. Nat. Rev. Mol. Cell Biol. 18, 407–422. 10.1038/nrm.2017.26.

23. Nodelman, I.M., and Bowman, G.D. (2021). Biophysics of Chromatin Remodeling. Annu. Rev. Biophys. 50, 73–93. 10.1146/annurev-biophys-082520-080201.

24. Larson, A.G., and Narlikar, G.J. (2018). The Role of Phase Separation in Heterochromatin Formation, Function, and Regulation. Biochemistry 57, 2540–2548. 10.1021/acs.biochem.8b00401.

25. Sabari, B.R., Dall’Agnese, A., and Young, R.A. (2020). Biomolecular Condensates in the Nucleus. Trends Biochem. Sci. 45, 961–977. 10.1016/j.tibs.2020.06.007.

26. Lyon, A.S., Peeples, W.B., and Rosen, M.K. (2021). A framework for understanding the functions of biomolecular condensates across scales. Nat. Rev. Mol. Cell Biol. 22, 215–235. 10.1038/s41580-020-00303-z.

27. Lafontaine, D.L.J., Riback, J.A., Bascetin, R., and Brangwynne, C.P. (2021). The nucleolus as a multiphase liquid condensate. Nat. Rev. Mol. Cell Biol. 22, 165–182. 10.1038/s41580-020-0272-6.

28. Mehta, S., and Zhang, J. (2022). Liquid–liquid phase separation drives cellular function and dysfunction in cancer. Nat. Rev. Cancer 22, 239–252. 10.1038/s41568-022-00444-7.

29. Holehouse, A.S., and Kragelund, B.B. (2023). The molecular basis for cellular function of intrinsically disordered protein regions. Nat. Rev. Mol. Cell Biol. 10.1038/s41580-023-00673-0.

30. Sabari, B.R., Hyman, A.A., and Hnisz, D. (2024). Functional specificity in biomolecular condensates revealed by genetic complementation. Nat. Rev. Genet. 10.1038/s41576-024-00780-4.

31. Pei, G., Lyons, H., Li, P., and Sabari, B.R. (2024). Transcription regulation by biomolecular condensates. Nat. Rev. Mol. Cell Biol. 10.1038/s41580-024-00789-x.

32. Cheng, Y., Shen, Z., Gao, Y., Chen, F., Xu, H., Mo, Q., Chu, X., Peng, C., McKenzie, T.T., Palacios, B.E., et al. (2022). Phase transition and remodeling complex assembly are important for SS18-SSX oncogenic activity in synovial sarcomas. Nat. Commun. 13, 2724. 10.1038/s41467-022-30447-9.

33. Patil, A., Strom, A.R., Paulo, J.A., Collings, C.K., Ruff, K.M., Shinn, M.K., Sankar, A., Cervantes, K.S., Wauer, T., Laurent, J.D.S., et al. (2023). A disordered region controls cBAF activity via condensation and partner recruitment. Cell 186, 4936-4955.e26. 10.1016/j.cell.2023.08.032.

34. Kim, Y.R., Joo, J., Lee, H.J., Kim, C., Park, J.-C., Yu, Y.S., Kim, C.R., Lee, D.H., Cha, J., Kwon, H., et al. (2024). Prion-like domain mediated phase separation of ARID1A promotes oncogenic potential of Ewing’s sarcoma. Nat. Commun. 15, 6569. 10.1038/s41467-024-51050-0.

35. Li, P., Zhai, Z., Fan, Y., Li, W., Ke, M., Li, X., Gao, H., Fu, Y., Ma, Z., Zhang, W., et al. (2024). Condensate remodeling reorganizes innate SS18 in synovial sarcomagenesis. Oncogenesis 13, 1–9. 10.1038/s41389-024-00539-w.

36. Davis, R.B., Supakar, A., Ranganath, A.K., Moosa, M.M., and Banerjee, P.R. (2024). Heterotypic interactions can drive selective co-condensation of prion-like low-complexity domains of FET proteins and mammalian SWI/SNF complex. Nat. Commun. 15, 1168. 10.1038/s41467-024-44945-5.

37. Tatavosian, R., Kent, S., Brown, K., Yao, T., Duc, H.N., Huynh, T.N., Zhen, C.Y., Ma, B., Wang, H., and Ren, X. (2019). Nuclear condensates of the Polycomb protein chromobox 2 (CBX2) assemble through phase separation. J. Biol. Chem. 294, 1451–1463. 10.1074/jbc.RA118.006620.

38. Plys, A.J., Davis, C.P., Kim, J., Rizki, G., Keenen, M.M., Marr, S.K., and Kingston, R.E. (2019). Phase separation of Polycomb-repressive complex 1 is governed by a charged disordered region of CBX2. Genes Dev. 33, 799–813. 10.1101/gad.326488.119.

39. Seif, E., Kang, J.J., Sasseville, C., Senkovich, O., Kaltashov, A., Boulier, E.L., Kapur, I., Kim, C.A., and Francis, N.J. (2020). Phase separation by the polyhomeotic sterile alpha motif compartmentalizes Polycomb Group proteins and enhances their activity. Nat. Commun. 11, 5609. 10.1038/s41467-020-19435-z.

40. Niekamp, S., Marr, S.K., Oei, T.A., Subramanian, R., and Kingston, R.E. (2024). Modularity of PRC1 composition and chromatin interaction define condensate properties. Mol. Cell 84, 1651-1666.e12. 10.1016/j.molcel.2024.03.001.

41. Gemeinhardt, T.M., Regy, R.M., Phan, T.M., Pal, N., Sharma, J., Senkovich, O., Mendiola, A.J., Ledterman, H.J., Henrickson, A., Lopes, D., et al. (2025). How a disordered linker in the Polycomb protein Polyhomeotic tunes phase separation and oligomerization. Preprint at bioRxiv, 10.1101/2023.10.26.564264 10.1101/2023.10.26.564264.

42. Gibson, B.A., Doolittle, L.K., Schneider, M.W.G., Jensen, L.E., Gamarra, N., Henry, L., Gerlich, D.W., Redding, S., and Rosen, M.K. (2019). Organization of Chromatin by Intrinsic and Regulated Phase Separation. Cell 179, 470-484.e21. 10.1016/j.cell.2019.08.037.

43. Schneider, M.W.G., Gibson, B.A., Otsuka, S., Spicer, M.F.D., Petrovic, M., Blaukopf, C., Langer, C.C.H., Batty, P., Nagaraju, T., Doolittle, L.K., et al. (2022). A mitotic chromatin phase transition prevents perforation by microtubules. Nature 609, 183–190. 10.1038/s41586-022-05027-y.

44. Moore, C., Wong, E., Kaur, U., Chio, U.S., Zhou, Z., Ostrowski, M., Wu, K., Irkliyenko, I., Wang, S., Ramani, V., et al. (2024). ATP-dependent remodeling of chromatin condensates uncovers distinct mesoscale effects of two remodelers. Preprint at bioRxiv, 10.1101/2024.09.10.611504 10.1101/2024.09.10.611504.

45. Grau, D.J., Chapman, B.A., Garlick, J.D., Borowsky, M., Francis, N.J., and Kingston, R.E. (2011). Compaction of chromatin by diverse Polycomb group proteins requires localized regions of high charge. Genes Dev. 25, 2210–2221. 10.1101/gad.17288211.

46. Isono, K., Endo, T.A., Ku, M., Yamada, D., Suzuki, R., Sharif, J., Ishikura, T., Toyoda, T., Bernstein, B.E., and Koseki, H. (2013). SAM Domain Polymerization Links Subnuclear Clustering of PRC1 to Gene Silencing. Dev. Cell 26, 565–577. 10.1016/j.devcel.2013.08.016.

47. Lau, M.S., Schwartz, M.G., Kundu, S., Savol, A.J., Wang, P.I., Marr, S.K., Grau, D.J., Schorderet, P., Sadreyev, R.I., Tabin, C.J., et al. (2017). Mutation of a nucleosome compaction region disrupts Polycomb-mediated axial patterning. Science 355, 1081–1084. 10.1126/science.aah5403.

48. Coré, N., Bel, S., Gaunt, S.J., Aurrand-Lions, M., Pearce, J., Fisher, A., and Djabali, M. Altered cellular proliferation and mesoderm patterning in Polycomb-M33-deficient mice.

49. Kundu, S., Ji, F., Sunwoo, H., Jain, G., Lee, J.T., Sadreyev, R.I., Dekker, J., and Kingston, R.E. (2017). Polycomb Repressive Complex 1 Generates Discrete Compacted Domains that Change during Differentiation. Mol. Cell 65, 432-446.e5. 10.1016/j.molcel.2017.01.009.

50. Wani, A.H., Boettiger, A.N., Schorderet, P., Ergun, A., Münger, C., Sadreyev, R.I., Zhuang, X., Kingston, R.E., and Francis, N.J. (2016). Chromatin topology is coupled to Polycomb group protein subnuclear organization. Nat. Commun. 7, 10291. 10.1038/ncomms10291.

51. Shao, Z., Raible, F., Mollaaghababa, R., Guyon, J.R., Wu, C., Bender, W., and Kingston, R.E. (1999). Stabilization of Chromatin Structure by PRC1, a Polycomb Complex. Cell 98, 37–46. 10.1016/S0092-8674(00)80604-2.

52. Francis, N.J., Saurin, A.J., Shao, Z., and Kingston, R.E. (2001). Reconstitution of a Functional Core Polycomb Repressive Complex. Mol. Cell 8, 545–556. 10.1016/S1097-2765(01)00316-1.

53. Stanton, B.Z., Hodges, C., Calarco, J.P., Braun, S.M.G., Ku, W.L., Kadoch, C., Zhao, K., and Crabtree, G.R. (2017). Smarca4 ATPase mutations disrupt direct eviction of PRC1 from chromatin. Nat. Genet. 49, 282–288. 10.1038/ng.3735.

54. Kadoch, C., Williams, R.T., Calarco, J.P., Miller, E.L., Weber, C.M., Braun, S.M.G., Pulice, J.L., Chory, E.J., and Crabtree, G.R. (2017). Dynamics of BAF–Polycomb complex opposition on heterochromatin in normal and oncogenic states. Nat. Genet. 49, 213–222. 10.1038/ng.3734.

55. Weber, C.M., Hafner, A., Kirkland, J.G., Braun, S.M.G., Stanton, B.Z., Boettiger, A.N., and Crabtree, G.R. (2021). mSWI/SNF promotes Polycomb repression both directly and through genome-wide redistribution. Nat. Struct. Mol. Biol. 28, 501–511. 10.1038/s41594-021-00604-7.

56. Kelso, T.W.R., Porter, D.K., Amaral, M.L., Shokhirev, M.N., Benner, C., and Hargreaves, D.C. (2017). Chromatin accessibility underlies synthetic lethality of SWI/SNF subunits in ARID1A-mutant cancers. eLife 6, e30506. 10.7554/eLife.30506.

57. Chory, E.J., Kirkland, J.G., Chang, C.-Y., D’Andrea, V.D., Gourisankar, S., Dykhuizen, E.C., and Crabtree, G.R. (2020). Chemical Inhibitors of a Selective SWI/SNF Function Synergize with ATR Inhibition in Cancer Cell Killing. ACS Chem. Biol. 15, 1685–1696. 10.1021/acschembio.0c00312.

58. Schiavoni, F., Zuazua-Villar, P., Roumeliotis, T.I., Benstead-Hume, G., Pardo, M., Pearl, F.M.G., Choudhary, J.S., and Downs, J.A. (2022). Aneuploidy tolerance caused by BRG1 loss allows chromosome gains and recovery of fitness. Nat. Commun. 13, 1731. 10.1038/s41467-022-29420-3.

59. Los, G.V., Encell, L.P., McDougall, M.G., Hartzell, D.D., Karassina, N., Zimprich, C., Wood, M.G., Learish, R., Ohana, R.F., Urh, M., et al. (2008). HaloTag: A Novel Protein Labeling Technology for Cell Imaging and Protein Analysis. ACS Chem. Biol. 3, 373–382. 10.1021/cb800025k.

60. Durisic, N., Laparra-Cuervo, L., Sandoval-Álvarez, Á., Borbely, J.S., and Lakadamyali, M. (2014). Single-molecule evaluation of fluorescent protein photoactivation efficiency using an in vivo nanotemplate. Nat. Methods 11, 156–162. 10.1038/nmeth.2784.

61. Skene, P.J., and Henikoff, S. (2017). An efficient targeted nuclease strategy for high-resolution mapping of DNA binding sites. eLife 6, e21856. 10.7554/eLife.21856.

62. Kim, C.A., Gingery, M., Pilpa, R.M., and Bowie, J.U. (2002). The SAM domain of polyhomeotic forms a helical polymer. Nat. Struct. Biol. 9, 5.

63. Brown, K., Chew, P.Y., Ingersoll, S., Espinosa, J.R., Aguirre, A., Kutateladze, T., Guevara, R.C., and Ren, X. (2022). Principles of assembly and regulation of condensates of Polycomb repressive complex 1 through phase separation (Biochemistry) 10.1101/2022.12.26.521954.

64. Jaensch, E.S., Zhu, J., Cochrane, J.C., Marr, S.K., Oei, T.A., Damle, M., McCaslin, E.Z., and Kingston, R.E. (2021). A Polycomb domain found in committed cells impairs differentiation when introduced into PRC1 in pluripotent cells. Mol. Cell 81, 4677-4691.e8. 10.1016/j.molcel.2021.09.018.

65. Richmond, E., and Peterson, C.L. (1996). Functional analysis of the DNA-stimulated ATPase domain of yeast SWI2/SNF2. Nucleic Acids Res. 24, 3685–3692.

66. Sif, S., Saurin, A.J., Imbalzano, A.N., and Kingston, R.E. (2001). Purification and characterization of mSin3A-containing Brg1 and hBrm chromatin remodeling complexes. Genes Dev. 10.1101/gad.872801. 15, 603–618.

67. Bultman, S.J., Gebuhr, T.C., and Magnuson, T. (2005). A Brg1 mutation that uncouples ATPase activity from chromatin remodeling reveals an essential role for SWI/SNF-related complexes in β-globin expression and erythroid development. Genes Dev. 19, 2849–2861. 10.1101/gad.1364105.

68. Pan, J., McKenzie, Z.M., D’Avino, A.R., Mashtalir, N., Lareau, C.A., St. Pierre, R., Wang, L., Shilatifard, A., and Kadoch, C. (2019). The ATPase module of mammalian SWI/SNF family complexes mediates subcomplex identity and catalytic activity–independent genomic targeting. Nat. Genet. 51, 618–626. 10.1038/s41588-019-0363-5.

69. Hodges, H.C., Stanton, B.Z., Cermakova, K., Chang, C.-Y., Miller, E.L., Kirkland, J.G., Ku, W.L., Veverka, V., Zhao, K., and Crabtree, G.R. (2018). Dominant-negative SMARCA4 missense mutations alter the accessibility landscape of tissue-unrestricted enhancers. Nat. Struct. Mol. Biol. 25, 61–72. 10.1038/s41594-017-0007-3.

70. Kim, J.M., Visanpattanasin, P., Jou, V., Liu, S., Tang, X., Zheng, Q., Li, K.Y., Snedeker, J., Lavis, L.D., Lionnet, T., et al. (2021). Single-molecule imaging of chromatin remodelers reveals role of ATPase in promoting fast kinetics of target search and dissociation from chromatin. eLife 10, e69387. 10.7554/eLife.69387.

71. Tilly, B.C., Chalkley, G.E., van der Knaap, J.A., Moshkin, Y.M., Kan, T.W., Dekkers, D.H., Demmers, J.A., and Verrijzer, C.P. (2021). In vivo analysis reveals that ATP-hydrolysis couples remodeling to SWI/SNF release from chromatin. eLife 10, e69424. 10.7554/eLife.69424.

72. Martin, B.J.E., Ablondi, E.F., Goglia, C., Mimoso, C.A., Espinel-Cabrera, P.R., and Adelman, K. (2023). Global identification of SWI/SNF targets reveals compensation by EP400. Cell 186, 5290-5307.e26. 10.1016/j.cell.2023.10.006.

73. Sahu, R.K., Dhakshnamoorthy, J., Jain, S., Folco, H.D., Wheeler, D., and Grewal, S.I.S. (2024). Nucleosome remodeler exclusion by histone deacetylation enforces heterochromatic silencing and epigenetic inheritance. Mol. Cell 84, 3175-3191.e8. 10.1016/j.molcel.2024.07.006.

74. Holehouse, A.S., and Alberti, S. (2025). Molecular determinants of condensate composition. Mol. Cell 85, 290–308. 10.1016/j.molcel.2024.12.021.

75. Lyons, H., Veettil, R.T., Pradhan, P., Fornero, C., De La Cruz, N., Ito, K., Eppert, M., Roeder, R.G., and Sabari, B.R. (2023). Functional partitioning of transcriptional regulators by patterned charge blocks. Cell 186, 327-345.e28. 10.1016/j.cell.2022.12.013.

76. Zijlmans, D.W., Stelloo, S., Bax, D., Yordanov, Y., Toebosch, P., Raas, M.W.D., Verhelst, S., Lamers, L.A., Baltissen, M.P.A., Jansen, P.W.T.C., et al. (2025). PRC1 and PRC2 proximal interactome in mouse embryonic stem cells. Cell Rep. 44. 10.1016/j.celrep.2025.115362.

77. Dobrinić, P., Szczurek, A.T., and Klose, R.J. (2021). PRC1 drives Polycomb-mediated gene repression by controlling transcription initiation and burst frequency. Nat. Struct. Mol. Biol. 28, 811–824. 10.1038/s41594-021-00661-y.

78. Zhou, H., Xiong, Y., Liu, Z., Hou, S., and Zhou, T. (2021). Expression and prognostic significance of CBX2 in colorectal cancer: database mining for CBX family members in malignancies and vitro analyses. Cancer Cell Int. 21, 402. 10.1186/s12935-021-02106-4.

79. Wang, L., Ren, B., Zhuang, H., Zhong, Y., and Nan, Y. (2021). CBX2 Induces Glioma Cell Proliferation and Invasion Through the Akt/PI3K Pathway. Technol. Cancer Res. Treat. 20, 15330338211045831. 10.1177/15330338211045831.

80. Kadoch, C., and Crabtree, G.R. (2015). Mammalian SWI/SNF chromatin remodeling complexes and cancer: Mechanistic insights gained from human genomics. Sci. Adv. 1, e1500447. 10.1126/sciadv.1500447.

81. Weissmann, F., Petzold, G., VanderLinden, R., Huis in ‘t Veld, P.J., Brown, N.G., Lampert, F., Westermann, S., Stark, H., Schulman, B.A., and Peters, J.-M. (2016). biGBac enables rapid gene assembly for the expression of large multisubunit protein complexes. Proc. Natl. Acad. Sci. 113, E2564–E2569. 10.1073/pnas.1604935113.

82. Nabet, B., Roberts, J.M., Buckley, D.L., Paulk, J., Dastjerdi, S., Yang, A., Leggett, A.L., Erb, M.A., Lawlor, M.A., Souza, A., et al. (2018). The dTAG system for immediate and target-specific protein degradation. Nat. Chem. Biol. 14, 431–441. 10.1038/s41589-018-0021-8.

83. Tchasovnikarova, I.A., Marr, S.K., Damle, M., and Kingston, R.E. (2022). TRACE generates fluorescent human reporter cell lines to characterize epigenetic pathways. Mol. Cell 82, 479-491.e7. 10.1016/j.molcel.2021.11.035.

84. Schindelin, J., Arganda-Carreras, I., Frise, E., Kaynig, V., Longair, M., Pietzsch, T., Preibisch, S., Rueden, C., Saalfeld, S., Schmid, B., et al. (2012). Fiji: an open-source platform for biological-image analysis. Nat. Methods 9, 676–682. 10.1038/nmeth.2019.

85. Robinson, J.T., Thorvaldsdóttir, H., Winckler, W., Guttman, M., Lander, E.S., Getz, G., and Mesirov, J.P. (2011). Integrative genomics viewer. Nat. Biotechnol. 29, 24–26. 10.1038/nbt.1754.

86. Bolger, A.M., Lohse, M., and Usadel, B. (2014). Trimmomatic: a flexible trimmer for Illumina sequence data. Bioinformatics 30, 2114–2120. 10.1093/bioinformatics/btu170.

87. Ewels, P., Magnusson, M., Lundin, S., and Käller, M. (2016). MultiQC: summarize analysis results for multiple tools and samples in a single report. Bioinformatics 32, 3047–3048. 10.1093/bioinformatics/btw354.

88. Langmead, B., and Salzberg, S.L. (2012). Fast gapped-read alignment with Bowtie 2. Nat. Methods 9, 357–359. 10.1038/nmeth.1923.

89. Dobin, A., Davis, C.A., Schlesinger, F., Drenkow, J., Zaleski, C., Jha, S., Batut, P., Chaisson, M., and Gingeras, T.R. (2013). STAR: ultrafast universal RNA-seq aligner. Bioinformatics 29, 15–21. 10.1093/bioinformatics/bts635.

90. Li, H., Handsaker, B., Wysoker, A., Fennell, T., Ruan, J., Homer, N., Marth, G., Abecasis, G., Durbin, R., and 1000 Genome Project Data Processing Subgroup (2009). The Sequence Alignment/Map format and SAMtools. Bioinformatics 25, 2078–2079. 10.1093/bioinformatics/btp352.

91. Zhang, Y., Liu, T., Meyer, C.A., Eeckhoute, J., Johnson, D.S., Bernstein, B.E., Nusbaum, C., Myers, R.M., Brown, M., Li, W., et al. (2008). Model-based Analysis of ChIP-Seq (MACS). Genome Biol. 9, R137. 10.1186/gb-2008-9-9-r137.

92. Quinlan, A.R., and Hall, I.M. (2010). BEDTools: a flexible suite of utilities for comparing genomic features. Bioinformatics 26, 841–842. 10.1093/bioinformatics/btq033.

93. Ramírez, F., Ryan, D.P., Grüning, B., Bhardwaj, V., Kilpert, F., Richter, A.S., Heyne, S., Dündar, F., and Manke, T. (2016). deepTools2: a next generation web server for deep-sequencing data analysis. Nucleic Acids Res. 44, W160–W165. 10.1093/nar/gkw257.

94. Liao, Y., Smyth, G.K., and Shi, W. (2014). featureCounts: an efficient general purpose program for assigning sequence reads to genomic features. Bioinformatics 30, 923–930. 10.1093/bioinformatics/btt656.

95. Perez, G., Barber, G.P., Benet-Pages, A., Casper, J., Clawson, H., Diekhans, M., Fischer, C., Gonzalez, J.N., Hinrichs, A.S., Lee, C.M., et al. (2025). The UCSC Genome Browser database: 2025 update. Nucleic Acids Res. 53, D1243–D1249. 10.1093/nar/gkae974.

96. O’Leary, N.A., Wright, M.W., Brister, J.R., Ciufo, S., Haddad, D., McVeigh, R., Rajput, B., Robbertse, B., Smith-White, B., Ako-Adjei, D., et al. (2016). Reference sequence (RefSeq) database at NCBI: current status, taxonomic expansion, and functional annotation. Nucleic Acids Res. 44, D733–D745. 10.1093/nar/gkv1189.

97. Love, M.I., Huber, W., and Anders, S. (2014). Moderated estimation of fold change and dispersion for RNA-seq data with DESeq2. Genome Biol. 15, 550. 10.1186/s13059-014-0550-8.

98. Ritchie, M.E., Phipson, B., Wu, D., Hu, Y., Law, C.W., Shi, W., and Smyth, G.K. (2015). limma powers differential expression analyses for RNA-sequencing and microarray studies. Nucleic Acids Res. 43, e47. 10.1093/nar/gkv007.

99. Korotkevich, G., Sukhov, V., Budin, N., Shpak, B., Artyomov, M.N., and Sergushichev, A. (2021). Fast gene set enrichment analysis. Preprint at bioRxiv, 10.1101/060012 10.1101/060012.

100. Chen, Y., Chen, L., Lun, A.T.L., Baldoni, P.L., and Smyth, G.K. (2025). edgeR v4: powerful differential analysis of sequencing data with expanded functionality and improved support for small counts and larger datasets. Nucleic Acids Res. 53, gkaf018. 10.1093/nar/gkaf018.

101. Lawrence, M., Huber, W., Pagès, H., Aboyoun, P., Carlson, M., Gentleman, R., Morgan, M.T., and Carey, V.J. (2013). Software for Computing and Annotating Genomic Ranges. PLOS Comput. Biol. 9, e1003118. 10.1371/journal.pcbi.1003118.

102. Subramanian, A., Tamayo, P., Mootha, V.K., Mukherjee, S., Ebert, B.L., Gillette, M.A., Paulovich, A., Pomeroy, S.L., Golub, T.R., Lander, E.S., et al. (2005). Gene set enrichment analysis: A knowledge-based approach for interpreting genome-wide expression profiles. Proc. Natl. Acad. Sci. 102, 15545–15550. 10.1073/pnas.0506580102.

103. Dyer, P.N., Edayathumangalam, R.S., White, C.L., Bao, Y., Chakravarthy, S., Muthurajan, U.M., and Luger, K. (2003). Reconstitution of Nucleosome Core Particles from Recombinant Histones and DNA. In Methods in Enzymology (Elsevier), pp. 23–44. 10.1016/S0076-6879(03)75002-2.

104. Flaus, A. (2011). Principles and practice of nucleosome positioning in vitro. Front. Life Sci. 5, 5–27. 10.1080/21553769.2012.702667.

105. Utley, R.T., Ikeda, K., Grant, P.A., Côté, J., Steger, D.J., Eberharter, A., John, S., and Workman, J.L. (1998). Transcriptional activators direct histone acetyltransferase complexes to nucleosomes. Nature 394, 498–502. 10.1038/28886.

106. Luger, K., Rechsteiner, T.J., and Richmond, T.J. (1999). Preparation of nucleosome core particle from recombinant histones. In Methods in Enzymology (Elsevier), pp. 3–19. 10.1016/S0076-6879(99)04003-3.

107. Niekamp, S., Stuurman, N., and Vale, R.D. (2020). A 6-nm ultra-photostable DNA FluoroCube for fluorescence imaging. Nat. Methods 17, 437–441. 10.1038/s41592-020-0782-3.

108. Sakuma, T., Nakade, S., Sakane, Y., Suzuki, K.-I.T., and Yamamoto, T. (2016). MMEJ-assisted gene knock-in using TALENs and CRISPR-Cas9 with the PITCh systems. Nat. Protoc. 11, 118–133. 10.1038/nprot.2015.140.

109. Ran, F.A., Hsu, P.D., Wright, J., Agarwala, V., Scott, D.A., and Zhang, F. (2013). Genome engineering using the CRISPR-Cas9 system. Nat. Protoc. 8, 2281–2308. 10.1038/nprot.2013.143.

110. Grimm, J.B., Xie, L., Casler, J.C., Patel, R., Tkachuk, A.N., Falco, N., Choi, H., Lippincott-Schwartz, J., Brown, T.A., Glick, B.S., et al. (2021). A General Method to Improve Fluorophores Using Deuterated Auxochromes. JACS Au 1, 690–696. 10.1021/jacsau.1c00006.

