## Supplementary Information for "Mutual Antagonism Between PRC1 Condensates and SWI/SNF in Chromatin Regulation"

### Supplementary Figures

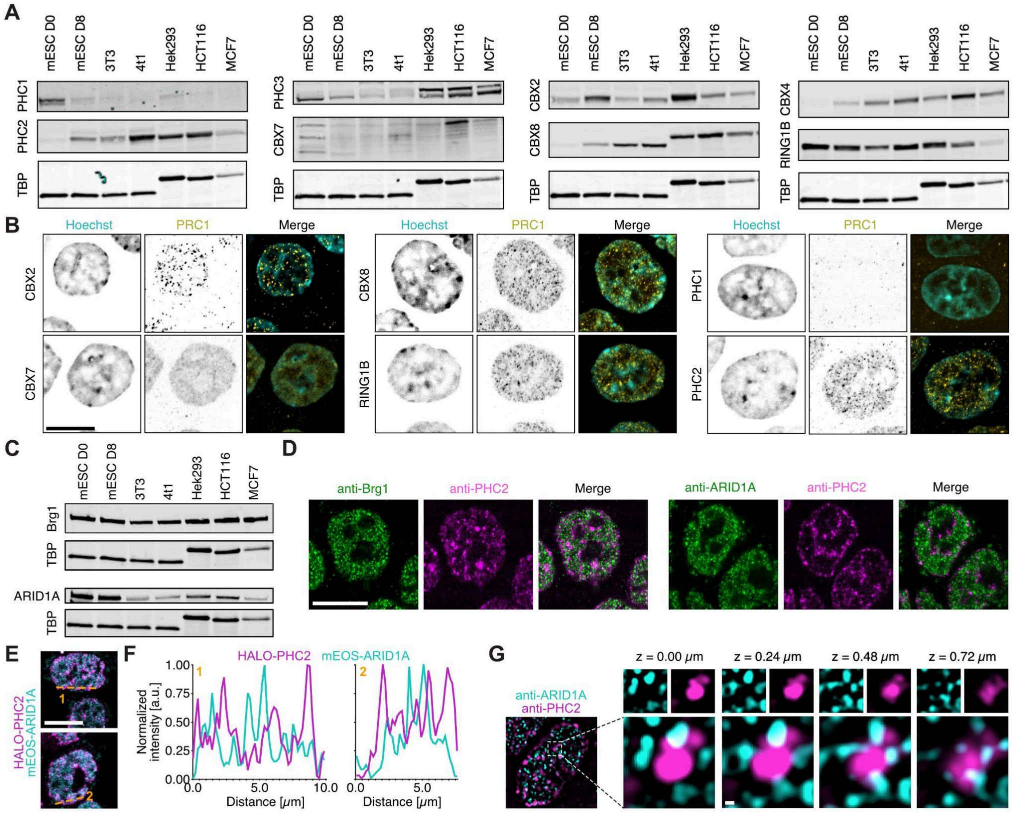


**Figure S1 | Characterization of HCT116 cell line for PRC1 and SWI/SNF subunits - related to Figure 1.** (**A**) Immunoblots of indicated PRC1 subunit expression in different cell lines: undifferentiated CJ7 mouse embryonic stem cells (mESC D0), mESCs differentiated to Neural Progenitors Cells (mESC D8), NIH-3T3 cells (3T3), 4T1 cells (4t1), Hek293 cells (Hek293), HCT116 cells (HCT116), and MCF7 cells (MCF7). Antisera against TATA-binding protein (TBP) was used as loading control. (**B**) Immunofluorescence of indicated PRC1 subunits in HCT116 cells. The scale bar is 10 μm. (**C**) Immunoblots of indicated SWI/SNF subunit expression in different cell lines: undifferentiated CJ7 mouse embryonic stem cells (mESC D0), mESCs differentiated to Neural Progenitors Cells (mESC D8), NIH-3T3 cells (3T3), 4T1 cells (4t1), Hek293 cells (Hek293), HCT116 cells (HCT116), and MCF7 cells (MCF7). Antisera against TATA-binding protein (TBP) was used as loading control. (**D**) Immunofluorescence of indicated SWI/SNF subunits in HCT116 cells. PHC2 was used as a reference. The scale bar is 10 μm. (**E**) Live-cell micrograph of endogenously-tagged PHC2 and ARID1A. The scale bar is 10 μm. To visualize endogenous SWI/SNF, we used the CRISPR-Cas9 based PITCh vector system[1,2] to insert mEOS3.2[3] at the N-terminus of ARID1A. Similarly, we tagged PHC2 with a HALO-tag[4] at its N-terminus, specifically labeling the long PHC2 isoform. The HALO-tag was then fluorescently labeled with HALO-JFX549[5]. (**F**) Line scan of intensities of data shown in **E** along the dashed lines. (**G**) Additional Structured Illumination Microscopy (SIM) micrographs of fixed HCT116 cells. Scale bar is 200 nm. Related to **Figure 1G**.

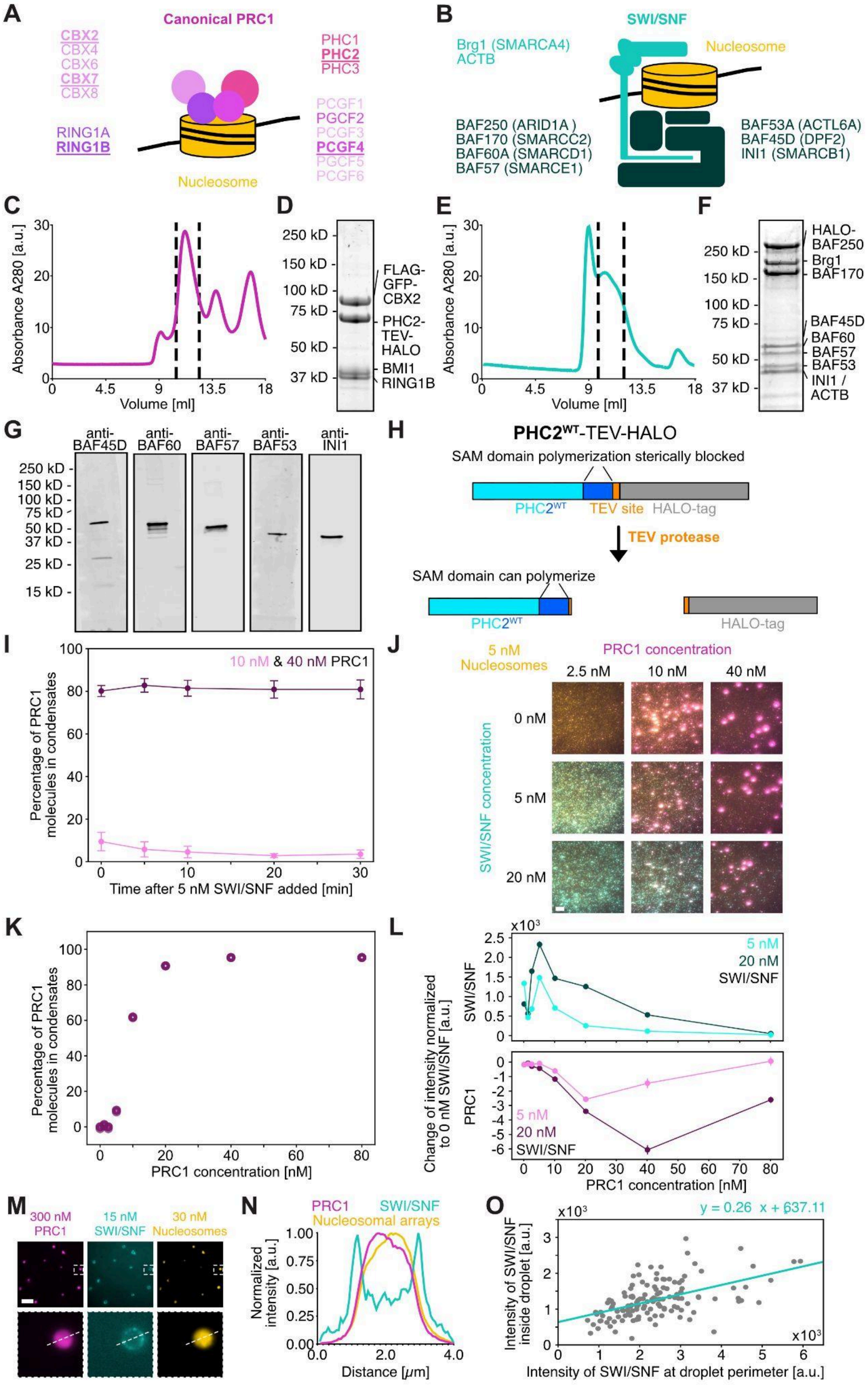


**Figure S2 | Purification of PRC1 and SWI/SNF complexes - related to Figure 2.** (**A**) Cartoon of canonical PRC1 composition. (**B**) Cartoon of SWI/SNF (cBAF) composition. (**C, E**) 280 nm absorbance traces from size exclusion chromatography for (**C**) PRC1 complexes with PHC2-TEV-HALO WT and (**E**) SWI/SNF. The area between the respective dashed lines indicate fractions that were pooled, concentrated, and analyzed by SDS-PAGE as shown in **D** and **F**. (**D**, **F**) 4-20% polyacrylamide gel of purified (**D**) canonical PRC1 complex with PHC2-TEV-HALO WT and (**F**) SWI/SNF. (**G**) Immunoblot of purified SWI/SNF complex staining for selected subunits. (**H**) Schematic showing how assays were performed with $PHC2^{WT}$ and $PHC2^{L307R}$. The addition of a cleavable polymerization blocker was necessary to enable purification of $PHC2^{WT}$ which would otherwise precipitate out as aggregates[6] (see **STAR Methods**). (**I**) Average percentage of PRC1 molecules in condensates over time - related to **Figure 2E-G**. (**J**) Example TIRF micrographs of nucleosomal arrays, SWI/SNF, and PRC1 mixed at various concentrations. Here, data was collected 30 minutes after SWI/SNF addition. The scale bar is 10 µm. (**K**) Percentage of PRC1 molecules in condensates as a function of PRC1 concentration. (**L**) Change of average PRC1 and SWI/SNF intensities on nucleosomal arrays as a function of various PRC1 concentrations. (**J-L**) Note, the data is partially the same as in **Figure 4B-D** (PRC1 first condition). For each condition 12,000 to >130,000 spots were analyzed. Data are representative of at least two technical replicates. (**M**) Representative epifluorescence microscopy micrographs when SWI/SNF is added to condensates of PRC1 and nucleosomal arrays. The scale bar is 10 µm. (**N**) Line scan of intensities of data shown in **M** along the dashed line. (**O**) Scatter plot of SWI/SNF intensity inside the condensate and at its periphery. 133 condensates were analyzed. Data are representative of three technical replicates.

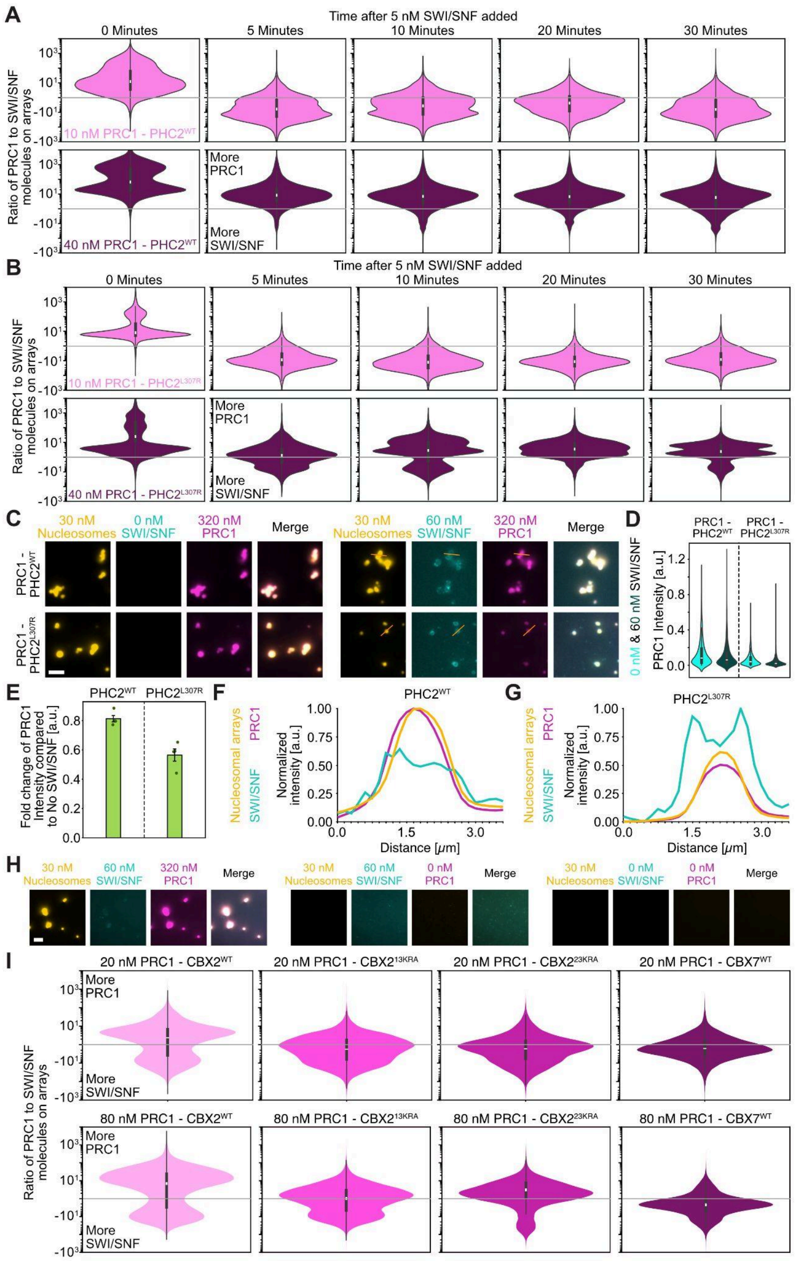


**Figure S3 | Role of PRC1 condensates on SWI/SNF binding to nucleosomal arrays - related to Figures 2 and 3.** (**A, B**) Violin plots of ratio of PRC1 to SWI/SNF molecules bound to nucleosomal arrays over time for (**A**) PHC2$^{WT}$ and (**B**) PHC2$^{L307R}$ - related to **Figure 2E-G**. Error bars are standard error of the mean of four technical replicates. For each condition 12,000 to >130,000 spots have been analyzed. (**C**) Representative epifluorescence microscopy micrographs when SWI/SNF is added to condensates of nucleosomal arrays and PRC1 with PHC2$^{WT}$ or PHC2$^{L307R}$. The scale bar is 5 μm. (**D**) Violin plots of PRC1 intensity of puncta of data as shown in **C**. (**E**) Bar graph of fold change of PRC1 intensity when SWI/SNF is added to condensates. Error bar is standard error of the mean. Note, the data is partially the same as in **Figure 4F**. (**F, G**) Line scan of intensities of data shown in **C** along the dashed line for PRC1 with either (**F**) $PHC2^{WT}$ or (**G**) $PHC2^{L307R}$. (**C-G**) Data are representative of four technical replicates. For each condition 1,400 to >11,000 spots were analyzed. (**H**) Representative Epi fluorescence microscopy micrographs when SWI/SNF is added to nucleosomal arrays and PRC1 (left) or to nucleosomal arrays alone (center) or when nucleosomal arrays are imaged alone (right). The scale bar is 5 μm. (**I**) Violin plots of ratio of PRC1 to SWI/SNF molecules bound to nucleosomal arrays for PRC1 with different CBX subunits incorporated - related to **Figure 3B-D**. For each condition 33,000 to >59,000 spots were analyzed. Data are representative of at least two technical replicates.

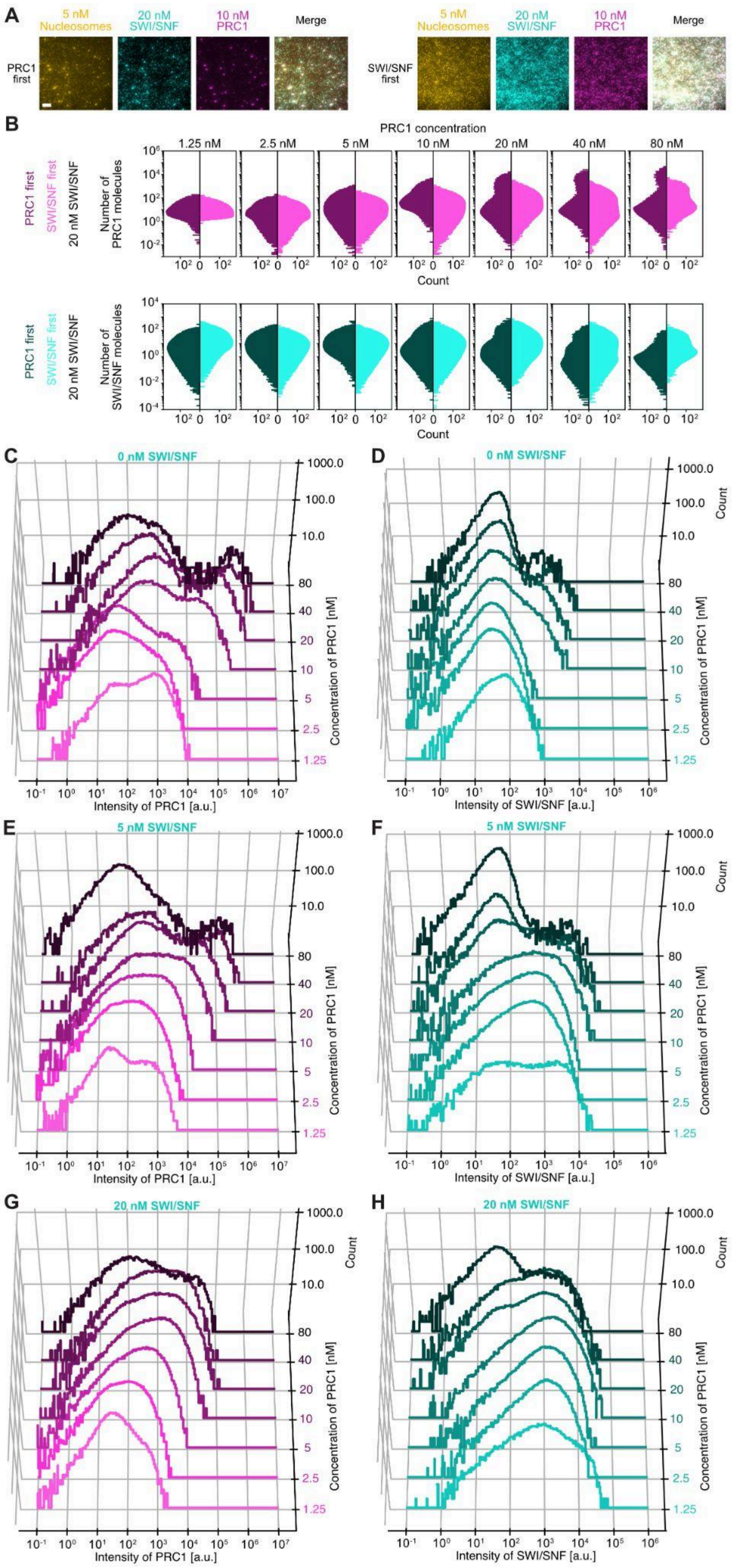


**Figure S4 | SWI/SNF presence on chromatin reduces PRC1 binding and condensate formation - related to Figure 4.** (**A**) Additional representative TIRF micrographs when

SWI/SNF and PRC1 are added to nucleosomal arrays in different orders. The scale bar is 10 μm - related to **Figure 4B**. (**B**) Intensity histograms of PRC1 and SWI/SNF for different order of addition - related to **Figure 4B-D**. (**C-H**) Histograms of PRC1 and SWI/SNF intensities when SWI/SNF is pre-incubated with nucleosomal arrays and PRC1 is added after 30 minutes - related to **Figure 4B-D**. (**B-H**) For each condition 29,000 to >105,000 spots were analyzed. Data are representative of at least two technical replicates.

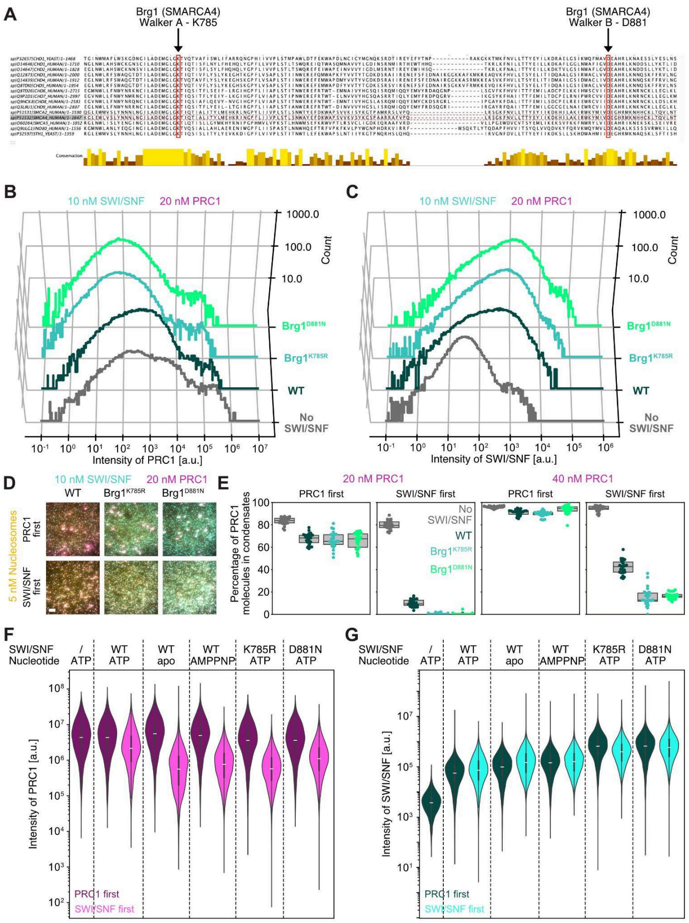


**Figure S5 | SWI/SNF presence on chromatin prevents PRC1 condensate formation in an ATP-hydrolysis independent manner - related to Figures 5.** (**A**) Sequence alignment of ATPase modules of different chromatin remodelers showing sequence conservation of Walker A and Walker B motifs. (**B, C**) Histograms of (**B**) PRC1 and (**C**) SWI/SNF intensities when nucleosomal arrays, PRC1, and SWI/SNF with different ATPase mutations are mixed (here PRC1 was added first) - related to **Figure 5C-F**. (**D**) Representative TIRF micrographs of nucleosomal arrays, PRC1, and different SWI/SNF complexes when SWI/SNF or PRC1 were added first. The scale bar is 10 µm. (**E**) Box plot of percentage of PRC1 molecules in condensates for data as shown in **D**. (**B-E**) Data are representative of at least two technical replicates. For each condition 44,000 to >98,000 spots were analyzed. (**F, G**) Violin plots of (**F**) PRC1 and (**G**) SWI/SNF intensities from epifluorescence microscopy experiments where 300 nM PRC1 with $PHC2^{WT}$, 15 nM SWI/SNF, and nucleosomal arrays (30 nM nucleosomes) have been mixed at various orders - related to **Figure 5**. Data are representative of at least two technical replicates. For each condition 8,000 to >93,000 spots were analyzed.

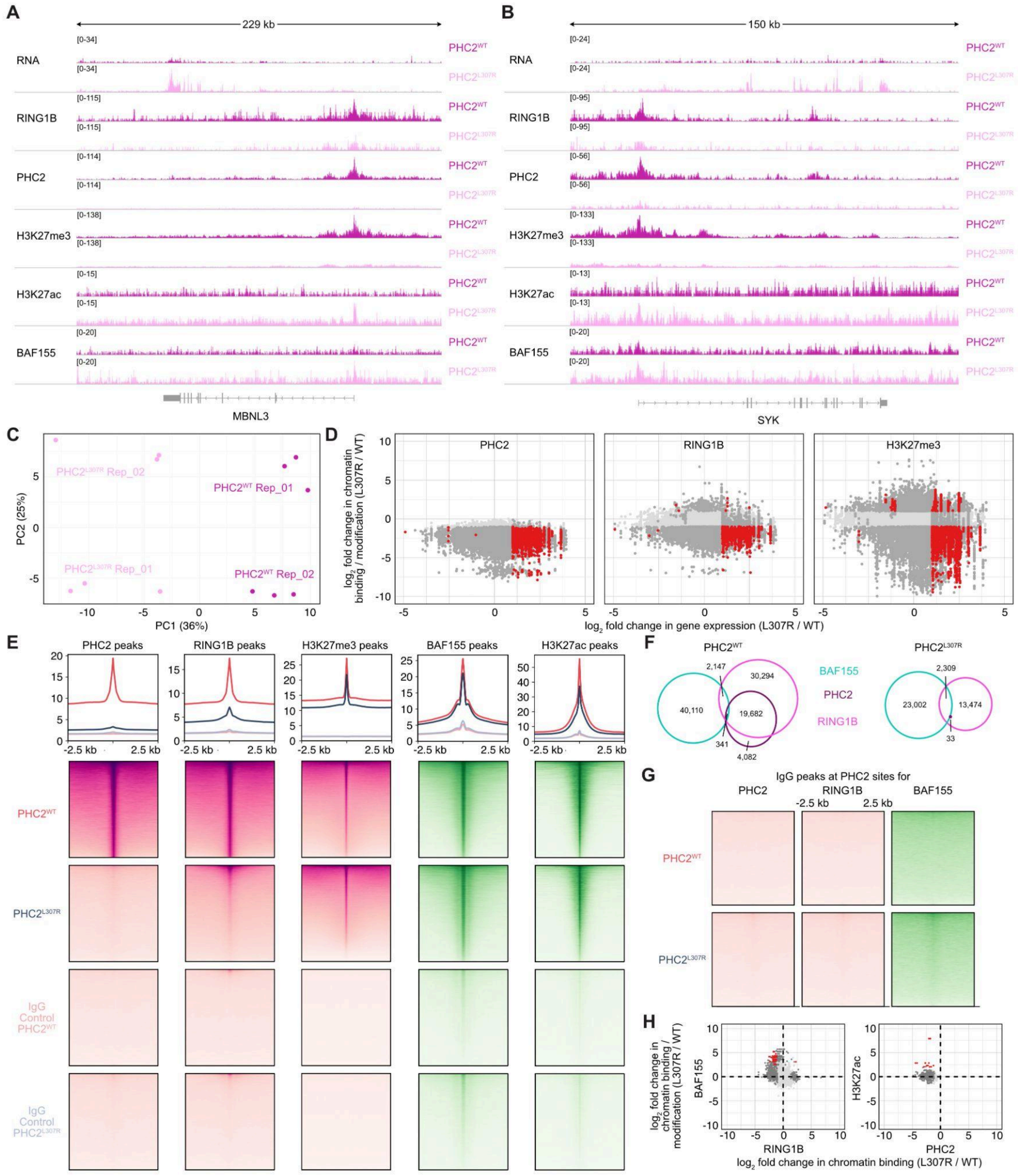


**Figure S6 | Loss of PHC2 polymerization derepresses genes and allows for SWI/SNF binding - related to Figure 6.** (**A, B**) Additional genome browser screenshots of RNAseq and CUT&RUN comparing HCT116 cells expressing endogenous PHC2$^{WT}$ or PHC2$^{L307R}$ - related to **Figure 6E**. (**C**) Principal component analysis of expression profiles of two different clones for

HCT116 cells expressing endogenous $PHC2^{WT}$ or $PHC2^{L307R}$ with three technical replicates each - related to **Figure 6F**. (**D**) Scatter plot correlating $log_2$ fold change in gene expression and chromatin binding for PHC2, RING1B, and H3K27me3 for HCT116 cells expressing endogenous $PHC2^{WT}$ or $PHC2^{L307R}$. Red dots are significant (log2 > 1; adj. P < 0.01) for both antibodies, dark grey for one antibody and light grey for neither antibody. Related to **Figure 6G**. (**E**) Heat maps showing CUT&RUN enrichment for PHC2, RING1B, H3K27me3, BAF155, and H3K27ac sites in HCT116 cells expressing endogenous $PHC2^{WT}$ or $PHC2^{L307R}$. IgG heatmaps are shown as control. (**F**) Venn diagrams of overlapping CUT&RUN peaks in HCT116 cells (log2 > 2; adj. P < 0.0001). (**G**) Heat maps showing CUT&RUN enrichment for IgG controls - related to **Figure 6I**. (**H**) Scatter plots of differential binding. Red dots are significant (log2 > 1; adj. P < 0.01) for both antibodies, dark grey for one antibody and light grey for neither antibody. (**A-H**) All experiments were performed with four to six replicates (two different clones and two to three technical replicates each).

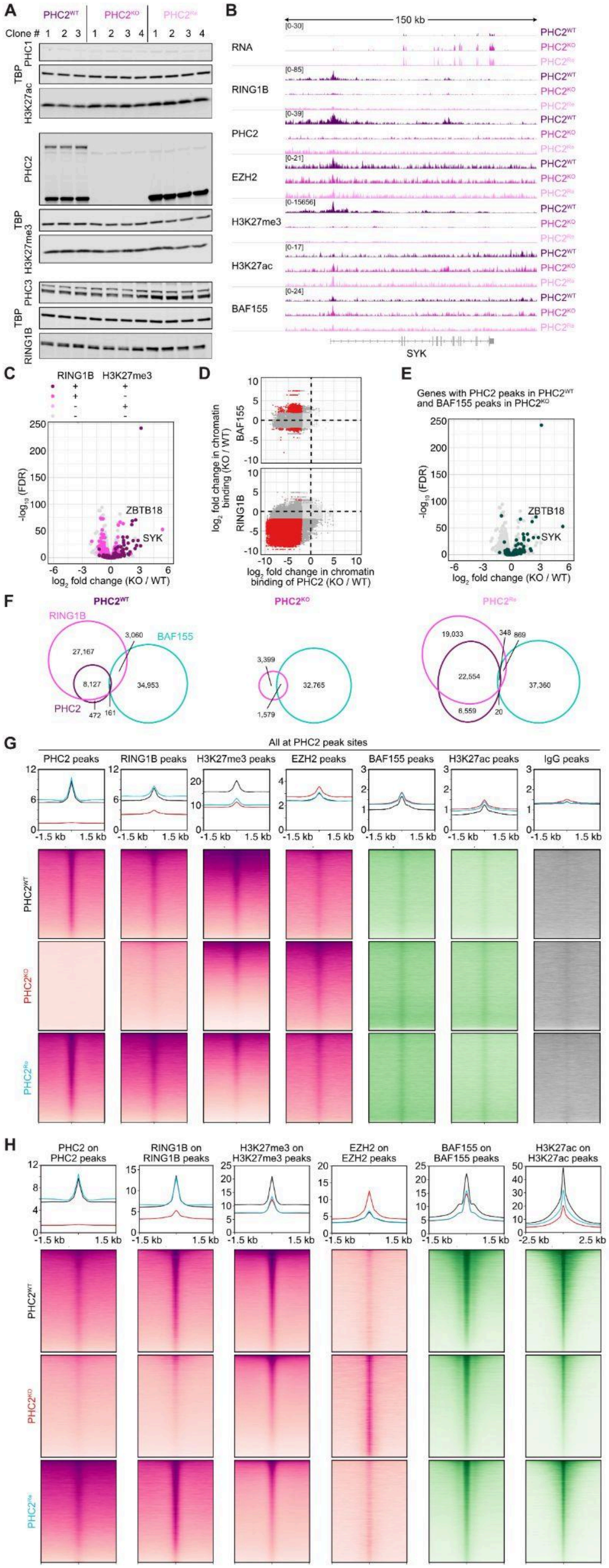
A
PHC2WT
PHC2KO
PHC2Re
Clone #
PHC1
TBP
H3K27ac
PHC2
H3K27me3
PHC3
RING1B
B
150 kb
RNA
RING1B
PHC2
EZH2
H3K27me3
H3K27ac
BAF155
SYK
C
RING1B
H3K27me3
ZBTB18
SYK
D
BAF155
RING1B
log2 fold change in chromatin binding (KO / WT)
log2 fold change in chromatin binding of PHC2 (KO / WT)
E
Genes with PHC2 peaks in PHC2WT and BAF155 peaks in PHC2KO
ZBTB18
SYK
-log10 (FDR)
log2 fold change (KO / WT)
F
RING1B
BAF155
PHC2
27,167
3,060
8,127
34,953
472
161
3,399
32,765
1,579
19,033
348
869
22,554
37,360
6,559
20
G
All at PHC2 peak sites
PHC2 peaks
RING1B peaks
H3K27me3 peaks
EZH2 peaks
BAF155 peaks
H3K27ac peaks
IgG peaks
-1.5 kb
1.5 kb
H
PHC2 on PHC2 peaks
RING1B on RING1B peaks
H3K27me3 on H3K27me3 peaks
EZH2 on EZH2 peaks
BAF155 on BAF155 peaks
H3K27ac on H3K27ac peaks
-2.5 kb
2.5 kb

**Figure S7 | SWI/SNF binding at lost PHC2 loci reduces PRC1 re-binding to chromatin - related to Figure 7.** (**A**) Immunoblot of HCT116 cells with wild type PHC2 ($PHC2^{WT}$), knockout of PHC2 ($PHC2^{KO}$), and re-expressed PHC2 ($PHC2^{Re}$). Antisera against TATA-binding protein (TBP) was used as loading control - related to **Figure 7A**. (**B**) Additional genome browser screenshot of RNAseq and CUT&RUN comparing HCT116 cells with $PHC2^{WT}$, $PHC2^{KO}$, or $PHC2^{Re}$ - related to **Figure 7E**. (**C**) Volcano plot reflecting changes in gene expression comparing HCT116 cells with $PHC2^{WT}$ or $PHC2^{KO}$. Pink and purple colored dots are genes reflecting changes in CUT&RUN peaks for RING1B and H3K27me3 (log2 > 1; adj. P < 0.01). (**D**) Scatter plots of differential binding in HCT116 cells with $PHC2^{WT}$ or $PHC2^{KO}$. Red dots are significant (log2 > 1; adj. P < 0.01) for both antibodies, dark grey for one antibody and light grey for neither antibody. Related to **Figure 7F**. (**E**) Volcano plot reflecting changes in gene expression comparing HCT116 cells expressing endogenous $PHC2^{WT}$ or $PHC2^{KO}$. Dark green colored dots are genes that have significant changes in CUT&RUN peaks for both, PHC2 in $PHC2^{WT}$ cells and for BAF155 in $PHC2^{KO}$ cells (log2 > 1; adj. P < 0.01). Related to **Figure 7F**. (**F**) Venn diagrams of overlapping CUT&RUN peaks in HCT116 cells with $PHC2^{WT}$, $PHC2^{KO}$, or $PHC2^{Re}$ (log2 > 2; adj. P < 0.0001). (**G**) Heat maps showing CUT&RUN enrichment at PHC2 peak sites for different antibodies of HCT116 cells with $PHC2^{WT}$, $PHC2^{KO}$, or $PHC2^{Re}$. (**H**) Heat maps showing CUT&RUN enrichment for PHC2, RING1B, H3K27me3, EZH2, BAF155, and H3K27ac sites in HCT116 cells with $PHC2^{WT}$, $PHC2^{KO}$, or $PHC2^{Re}$. (**B-H**) All experiments were performed with four replicates (two different clones and two technical replicates each).

#### Legends for Supplementary Movies

**Movie S1**

The movie shows Structured Illumination Microscopy (SIM) data of fixed HCT116 cells that were labeled with anti-PHC2 (pink) and anti-RING1B (green) antibodies (left) and anti-PHC2 (pink) and anti-ARID1A (green) antibodies (right). The scale bar is 5 μm.

**Movie S2**

The movie shows Structured Illumination Microscopy (SIM) data of fixed HCT116 cells that were labeled with anti-PHC2 (pink) and anti-ARID1A (cyan) antibodies as shown in **Figure 1D and 1E**. The scale bar is 200 nm.

**Movie S3**

The movie shows reconstituted PRC1 complexes with CBX2 and $PHC2^{WT}$ (pink), $SWI/SNF^{WT}$ (cyan), and nucleosomal array (yellow) after mixing all three as shown in **Figure 2B and 2C**. The duration of the movie is 28.5 minutes. The scale bar is 5 μm.

#### Supplementary References


1. Sakuma, T., Nakade, S., Sakane, Y., Suzuki, K.-I.T., and Yamamoto, T. (2016). MMEJ-assisted gene knock-in using TALENs and CRISPR-Cas9 with the PITCh systems. Nat. Protoc. *11*, 118–133. https://doi.org/10.1038/nprot.2015.140.

2. Nabet, B., Roberts, J.M., Buckley, D.L., Paulk, J., Dastjerdi, S., Yang, A., Leggett, A.L., Erb, M.A., Lawlor, M.A., Souza, A., et al. (2018). The dTAG system for immediate and target-specific protein degradation. Nat. Chem. Biol. *14*, 431–441. https://doi.org/10.1038/s41589-018-0021-8.

3. Durisic, N., Laparra-Cuervo, L., Sandoval-Álvarez, Á., Borbely, J.S., and Lakadamyali, M. (2014). Single-molecule evaluation of fluorescent protein photoactivation efficiency using an in vivo nanotemplate. Nat. Methods *11*, 156–162. https://doi.org/10.1038/nmeth.2784.

4. Los, G.V., Encell, L.P., McDougall, M.G., Hartzell, D.D., Karassina, N., Zimprich, C., Wood, M.G., Learish, R., Ohana, R.F., Urh, M., et al. (2008). HaloTag: A Novel Protein Labeling Technology for Cell Imaging and Protein Analysis. ACS Chem. Biol. *3*, 373–382. https://doi.org/10.1021/cb800025k.

5. Grimm, J.B., Xie, L., Casler, J.C., Patel, R., Tkachuk, A.N., Falco, N., Choi, H., Lippincott-Schwartz, J., Brown, T.A., Glick, B.S., et al. (2021). A General Method to Improve Fluorophores Using Deuterated Auxochromes. JACS Au *1*, 690–696. https://doi.org/10.1021/jacsau.1c00006.

6. Niekamp, S., Marr, S.K., Oei, T.A., Subramanian, R., and Kingston, R.E. (2024). Modularity of PRC1 composition and chromatin interaction define condensate properties. Mol. Cell *84*, 1651-1666.e12. https://doi.org/10.1016/j.molcel.2024.03.001.
